# ngsAMOVA: A Probabilistic Framework for Analysis of Molecular Variance, *d_XY_* and Neighbor-Joining Trees with Low Depth Sequencing Data

**DOI:** 10.1101/2025.05.12.653431

**Authors:** Isin Altinkaya, Lei Zhao, Rasmus Nielsen, Thorfinn Sand Korneliussen

## Abstract

**Motivation:** Next-generation sequencing (NGS) has transformed population genetics and evolutionary biology, but the data produced in studies of non-model organisms, ancient DNA, and environmental DNA often consist of low- or medium-depth sequencing. Analyses of these data rely on computational methods that utilize genotype likelihoods (GLs) to account for genotype uncertainty. Nevertheless, many widely-used analysis methods, such as analysis of molecular variance (AMOVA) and methods for estimating phylogenetic trees using nucleotide divergence (*dXY*) still lack the probabilistic frameworks necessary to accommodate GLs.

**Results:** We introduce ngsAMOVA, a novel probabilistic framework for analyzing molecular variation in population hierarchies with low- and medium-depth sequencing data. It employs an Expectation Maximization algorithm to first estimate the joint genotype probabilities for pairs of individuals, accounting for genotype uncertainty using GLs. It then uses these estimates to generate a pairwise distance matrix, which can be used for AMOVA, estimation of *dXY*, and for estimating phylogenetic trees using Neighbor-Joining. Hypothesis testing is facilitated using genomic block-bootstrapping. Through extensive simulations, we demonstrate that ngsAMOVA provides more accurate results compared to genotype calling at low and medium read depths. Overall, ngsAMOVA represents a methodological advance in the analysis of molecular variance and divergence under sequencing uncertainty. It provides a robust framework, opening up numerous possibilities for gaining insights into the evolutionary histories through its applications. ngsAMOVA is available as a fast, efficient, and user-friendly program written in C/C++.

**Availability:** ngsAMOVA is freely available at https://github.com/isinaltinkaya/ngsAMOVA.

**Supplementary information:** Supplementary data are available online.

## 1 Introduction

Next-generation sequencing (NGS) technologies have revolutionized evolutionary biology and molecular ecological research by facilitating genome-scale analysis across diverse contexts, including studies of non-model organisms, ancient DNA, and environmental DNA data (Knapp and Hofreiter, 2010; Shokralla *et al*., 2012; da Fonseca *et al*., 2016). Datasets obtained in such studies are often generated using low or medium sequencing depth, resulting in significant genotype uncertainty. Conventional methods for the analysis of molecular variance (AMOVA), estimation of absolute nucleotide divergence (*d*_*XY*_), and phylogenetic tree construction, rely on genotypes that are called with high confidence. This dependence presents substantial issues, as genotype calling at low depths may result in biased estimates and/or considerable loss of informative sites due to stringent filtering criteria.

AMOVA is a widely used method for identifying and quantifying genetic variance and population structures by decomposing genetic variance into hierarchical components (Excoffier *et al*., 1992). It provides an effective framework for hypothesis testing for hierarchical population structure and differentiation, primarily due to its high interpretability. A number of software tools exist for performing conventional AMOVA analyses, such as `Arlequin` (Excoffier *et al*., 2005), `poppr` (Kamvar *et al*., 2014), `ade4` (Thioulouse *et al*., 2018), `GenAlEx` (Peakall and Smouse, 2006, 2012), `GENODIVE` (Meirmans, 2020), `vcfpop` (Huang *et al*., 2022), `polygene` (Huang *et al*., 2020). Several extensions of the conventional AMOVA framework exist, such as a simulated annealing procedure aiming to maximize the proportion of total genetic variance due to differences between groups (Dupanloup *et al*., 2002), a generalized AMOVA framework (Nievergelt *et al*., 2007), an AMOVA-based clustering where the F-statistics from AMOVA are used as the optimality criterion to find the clustering that gives the maximum amount of genetic differentiation among clusters (Meirmans, 2012, 2020), a generalized AMOVA framework for any number of ploidies and hierarchies (Huang *et al*., 2021), and an AMOVA framework for autopolyploids (Meirmans and Liu, 2018). However, existing AMOVA frameworks do not address genotype uncertainty and their hypothesis testing methods are not designed for genomic data. This limitation can significantly affect its accuracy and reliability when applied to genomic datasets characterized by such uncertainty.

The absolute nucleotide divergence, *d*_*XY*_, measures the average genetic distance between two populations by calculating pairwise differences among individuals from the two populations (Nei and Li, 1979). Despite its frequent use for assessing population divergence, current implementations of *d*_*XY*_ similarly rely on accurately called genotypes and may be biased when applied to low-depth sequencing data.

Neighbor-Joining (NJ) is a popular distance-based phylogenetic method utilized for constructing simple evolutionary trees due to its computational simplicity and statistical consistency (Saitou and Nei, 1987). NJ uses distances genetic and may also be affected by genotype calling errors when applied to low-coverage NGS datasets. This underscores the need for developing new methods for estimating genetic distances that can incorporate GLs.

## 2 Materials and methods

Here, we develop a new framework for estimating *d*_*XY*_ and for AMOVA based on GLs. There are three steps in the inference framework. We first use an EM algorithm to obtain maximum likelihood estimates of the joint genotype probabilities for pairs of individuals, using GLs to account for genotype uncertainty. Second, we compute pairwise distances between individuals. Third, we use this distance matrix, along with the hierarchical grouping information associated with each individual pair, to perform AMOVA, estimate the absolute nucleotide divergence *d*_*XY*_, and construct phylogenetic trees using NJ. We implemented these methods using C++ and htslib to ensure fast and memory-efficient analyses (Bonfield *et al*., 2021) and we also implemented an option to use genotype data (`–bcf-src 2`). Measures of statistical confidence can be calculated using a block bootstrap approach.

### 2.1 The Expectation-Maximization Algorithm

For each pair of individuals, we define *M* to be the 3 *×* 3 matrix of joint genotype probabilities, with columns representing the genotype of one individual (e.g., 0, 1, or 2) and rows representing the genotypes of the other individual, assuming di-allelic data. We estimate *M* using an EM algorithm that iteratively updates the entries of *M* and terminates based on two criteria: (1) when the increase in likelihood between successive iterations falls below a predefined tolerance threshold, or (2) when the maximum allowable number of iterations is reached (a predefined iteration count threshold). The EM algorithm is described in detail in S1.3.

### 2.2 Pairwise Distance Matrix and *dxy*

We define the pairwise differences as the probability that two alleles, one sampled from each individual, are not identical to each other. This distances can be calculated from *M* by taking a weighted sum over the appropriate elements of *M* (S1.1), and corresponds to traditional IBD distances under the infinite sites model assumption. If the distance between individual *i* and *j* is *d*_*ij*_, *d*_*XY*_ is then calculated in the standard way as

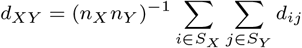

where *S*_*X*_ is the set of the *n*_*X*_ individuals from population *X* and *S*_*Y*_ is the set of the *n*_*Y*_ individuals from population *Y* (cite Nei and Li).

A key assumption in the AMOVA framework is that the input pairwise genetic distances are Euclidean distances. We therefore transform the genetic distance matrix into a Euclidean form using the Cailliez transformation (Cailliez, 1983) for the purpose of AMOVA analyses (S1.1).

### 2.3 AMOVA and Block Bootstrapping Framework

AMOVA assumes a fixed number, *L*, of hierarchical components and represents the total molecular variance 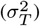 as the sum of its hierarchical covariance components, 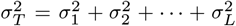. For example, for *L* = 3, we can decompose the total genetic variance into 3 components: among regions 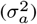, among populations within regions 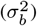, and among individuals within populations 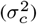, where 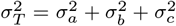 (S2.8). The AMOVA Φ*−* statistic is then calculated as

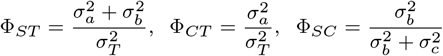

with a higher Φ-statistic value indicating a higher amount of differentiation. Here, F_*ST*_ represents the variance within populations and regions compared to the total, F_*CT*_ represents the variance among populations compared to the total, and F_*SC*_ represents the variance among populations compared to that within regions. This provides a framework for comparing the proportion of genetic variance across hierarchical levels.

The original AMOVA framework uses a permutation test to quantify the significance of Φ-statistics (Excoffier *et al*., 1992). This approach involves permuting units at a given hierarchical level and recalculating the Φ-statistic for each permuted dataset. By repeating this procedure many times, a distribution of the Φ-statistics under the null hypothesis that there is no genetic stratification with regard to the corresponding hypothesis, is obtained. However, this approach is limited as the accuracy of the significance test heavily depends on the number of possible ways to allocate the hierarchical components in test. Moreover, it has been shown that a hierarchical structure must have at least 6 groups to allow obtaining P-values smaller than 0.05 (Fitzpatrick, 2009). In genomic data containing substantial information from many independent genomic regions, the traditional approach of permuting components will necessarily be underpowered. To address this issue, we propose an alternative approach based on block bootstrapping.

To estimate the sampling variance of the Φ-statistic, we employ a block bootstrapping method, a resampling approach commonly utilized in genomic studies to account for linkage disequilibrium (LD) and to construct confidence intervals **???**. Specifically, we generate a distribution of resampled Φ-statistics, denoted F^*^, by repeatedly computing Φ on datasets constructed from genomic blocks sampled with replacement while preserving the correlation structure among individuals (S1). This approach preserves the local correlation structure among SNPs within the genomic blocks and provides an empirical estimate of the variance of Φ. To construct confidence intervals, the method implements the Bootstrap Percentile Method, Basic Percentile Method and Normal Approximation Method for constructing nonparametric confidence intervals for the original Φ estimate (S2.10.5).

An advantage of the Block Bootstrap procedure is that it accommodates LD as long as the block sizes are sufficiently large. We present a custom R script `get_block_size.R` for estimating the block size parameter based on LD decay estimated by `ngsLD` (Fox *et al*., 2019). `ngsLD` is first used to estimate the pairwise LD between sites using GLs, and then fit a decay model to the results. We modified the LD decay model fit script from `ngsLD` to produce a block size estimate based on the percentage decline of LD (S2.10.1). This block size can be supplied as a parameter to construct artificial genomic intervals.

### 2.4 Benchmarking

To evaluate the accuracy and robustness of our method, we simulated data under varying scenarios of population structure, sequencing depth and contig sizes. We used `msprime` (Baumdicker *et al*., 2022) to generate 20 replicates from three coalescent models, each simulating four structured populations (popA, popB, popC, popD) organized into two regions (reg1 and reg2). In all scenarios, we sampled 10 diploid individuals from each population, yielding a total of 40 individuals per replicate.

The demographic models were specified using the `demes` format (Gower *et al*., 2022) (S3.1) and visualized with `demesdraw` (Gower *et al*., 2022) (Figures S4, and S5, S6). Model 1 represents a scenario of regional isolation, allowing gene flow only between populations within the same region. Model 2 simulates reduced regional differentiation by introducing migration between populations across regions. Model 3 depicts complete isolation following population and regional splits, with no gene flow occurring post-divergence. All simulations used fixed recombination and mutation rates of 1.14856*×*10^−8^ and 1.29*×*10^−8^ per site per generation, respectively.

Using the ground-truth genomes from coalescence simulations, we generated low-to high-depth sequencing data with a fixed base-calling error rate of 0.2% using `vcfgl` (Altinkaya *et al*., 2025). Sequencing depths ranged from ultra-low to high coverage (0.01, 0.1, 0.5, 1, 2, 5, 10, and 100). These simulations included both variable and invariant sites to reflect realistic genomic data structures (S3.2). For genotype calling, we used the naive genotype calling method implemented in `BCFtools`, which selects the most likely genotype for each site (`–GL-to-GT` with `–threshold 1`) using the simulated GLs (S3.3).

## 3 Results and discussion

To evaluate the performance of our method, we used ngsAMOVA both with GL data and genotype calls to estimate AMOVA Φ-statistics, *d*_*XY*_, and construct NJ trees. The results were compared among models and depths to assess accuracy and robustness to sequencing uncertainty. Comparisons across demographic models, sequencing depths, and contig sizes consistently revealed that the GL-based approach outperformed genotype-based inference in terms of accuracy and robustness (Figures 1, 2, and 3).

**Fig. 1.**
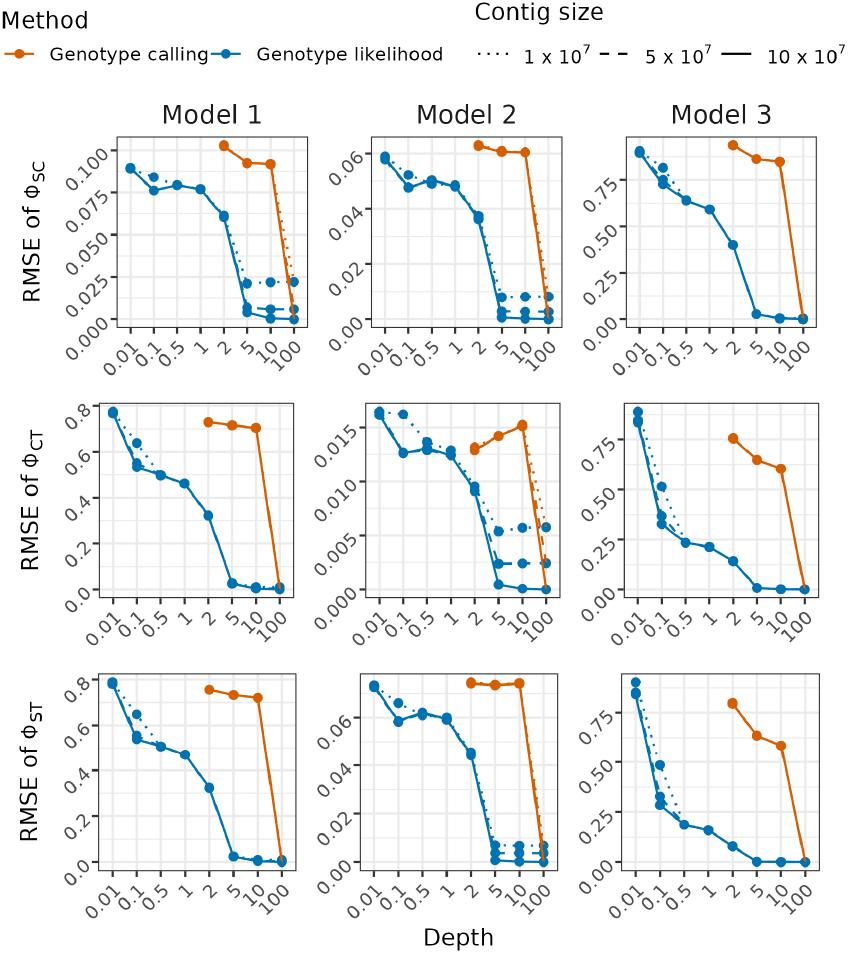
Root Mean Squared Error (RMSE) of Φ-statistic estimates across sequencing depths for different models and inference methods. Each row corresponds to a different AMOVA Φ-statistic: F_*SC*_ for population in region (top), F_*CT*_ for region in total (middle), and F_*ST*_ for population in total (bottom). Columns represent three models. The x-axis shows sequencing depth, and the y-axis shows the RMSE of each Φ-statistic, calculated in comparison to those obtained from ground-truth data for the contig size 1 × 10^7^. Two inference methods are compared: genotype calling (orange) and GL (blue), across three contig sizes (indicated by line types: dotted for 1 × 10^7^, dashed for 5 × 10^7^, and solid for 1 × 10^8^).

**Fig. 2.**
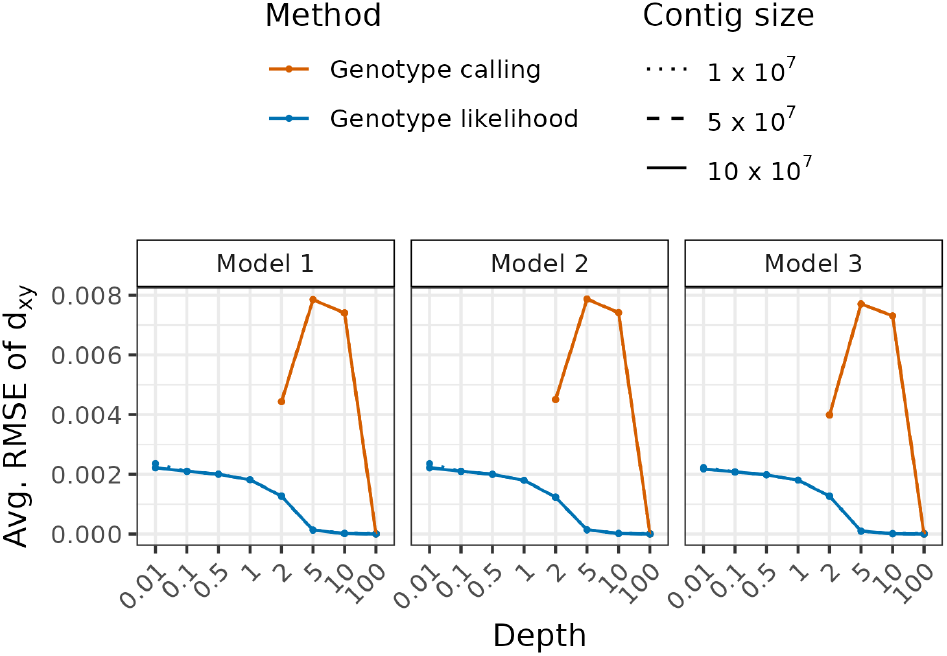
RMSE of *d*_*XY*_ estimates across varying sequencing depths (x-axis) under the three demographic models (columns), averaged across all *d*_*XY*_ comparisons (reg1 vs. reg2, popA vs. popB, popA vs. popC, popB vs. popC, popA vs. popD, popB vs. popD, and popC vs. popD). Results are indicated by colors for two methods: genotype calling (orange) and genotype likelihood (blue). Line styles indicate contig sizes: 1 × 10^7^ (dotted), 5 × 10^7^ (dashed), and 10 × 10^7^ (solid).

**Fig. 3.**
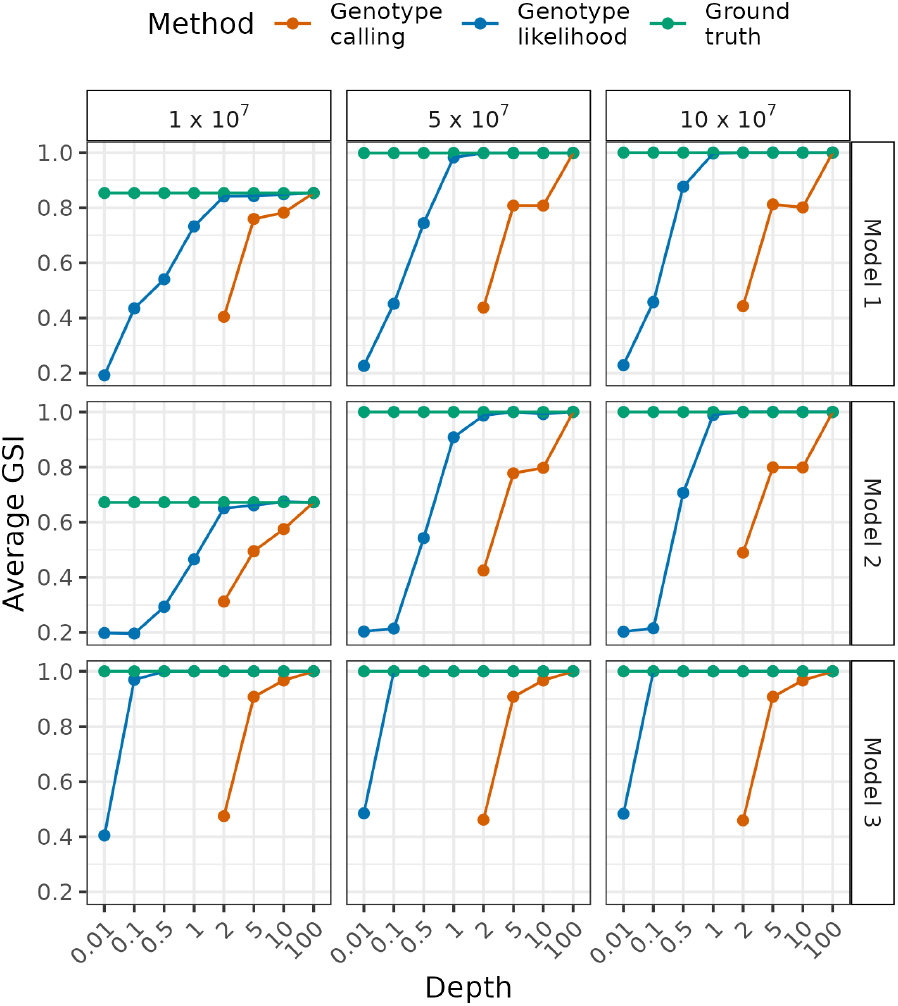
The Average Genealogical Sorting Index (GSI) values for different read depths, evaluated under three demographic models (rows: Model 1, Model 2, Model 3) and three contig sizes (columns: 1 × 10^7^, 5 × 10^7^, 10 × 10^7^ base pairs). Each point represents the mean GSI across 20 replicates for a given combination of depth and model. Methods compared include GL-based inference (blue), genotype calling (orange), and ground-truth based on ground-truth genotypes (green).

Benchmarking based on the absolute error of the pairwise distance estimates showed that the GL method consistently produced more accurate and stable results across individual pairs and replicates at all simulated sequencing depths below 100*×*, compared to the genotype calling approach (Figure S7). The accuracy of pairwise distance estimation remained high even at sequencing depths as low as 0.01*×*, with error rates declining steadily as depth increases.

To assess the accuracy of F-statistic estimates under varying conditions, we calculated the root mean square error (RMSE) of F_*SC*_, F_*CT*_, and F_*ST*_ for varying sequencing depths, contig sizes, and demographic models (Figure 1). Across all three models and contig sizes, the GL-based method consistently produced lower RMSE values compared to genotype calling, while genotype calling yielded high error rates even at 5*×* and 10*×* coverage. The GL-based approach showed a decline in RMSE with increasing depth and achieved near-zero error starting at 5*×* depth, highlighting the robustness of the genotype likelihood framework in accurately quantifying genetic structure and divergence, even under challenging sequencing conditions.

The average RMSE of the *d*_*XY*_ estimates for different estimation methods and models at different depths, averaged across all *d*_*XY*_ comparisons, are shown in Figure 2. The RMSE of *d*_*XY*_ for all different scenarios, methods, and models at different depths is shown in Figure S8. The GL results indicate a stable RMSE around 0.002 for depths *≤* 1, while the genotype calling results show a higher RMSE, indicating that the GL method captures inter-population divergence with higher greater accuracy than genotype-based alternatives across different depths and *d*_*XY*_ comparisons (Figure S8).

Our results also show that ngsAMOVA produces superior phylogenetic reconstructions, as evidenced by higher Genealogical Sorting Index (GSI) (Cummings *et al*., 2008) values and lower proportions of non-monophyletic populations across depths and contig sizes (Figure 3). The GSI is a measure of the degree of exclusive ancestry of labeled populations in the tree, and the ratio of non-monophyletic populations is the proportion of populations that are not monophyletic in the tree. The GL-based method produced tree estimates with a higher degree of exclusive ancestry compared to the genotype calling method. The results obtained using the GL-based method at 0.1*×*, and for some scenarios, 0.01*×* depth were comparable to those obtained using genotypes at 2*×* depth, below which the genotype calling method failed to construct trees due to missing pairwise distances. Additionally, average GSI values derived from ground-truth genotypes using smaller contig sizes remained below 1 in two of the simulated scenarios, indicating that the number of informative sites was insufficient to reconstruct trees that always group individuals with their populations, even when using the ground-truth genotypes. The average ratio of non-monophyletic populations results supported the findings from the average GSI results, showing that the GL method provided tree estimates with a lower ratio of non-monophyletic populations compared to the genotype calling method (Figure S9). This suggests that, given sufficient number of informative sites, the probabilistic treatment of genotype uncertainty preserves sufficient signal to accurately reconstruct population-level relationships.

Our results also showed the importance of the number of sites used in the analysis for obtaining accurate results. Specifically, we showed that increasing the number of sites allowed us to encapsulate more information, which led to more reliable estimates. When using standard genotype calling pipelines, strict thresholds are typically applied to ensure the reliability of calls. This can greatly reduce the number of sites available for analysis, which can have a drastic effect on the accuracy of the results. This underlines the importance of carefully considering the trade-off between the confidence in the called genotypes and the number of sites retained that would be available for downstream analysis. This limitation does not apply to the GL-based approach, as genotype likelihoods allow the inclusion of all sites. By accounting for uncertainty rather than filtering uncertain genotypes, the GL method retains more sites for analysis, thereby increasing the amount of information available and improving the reliability of the resulting estimates.

The results of our benchmarking analyses highlighted the limitations of traditional genotype-based methods when applied to low-coverage sequencing data, and demonstrate the effectiveness of ngsAMOVA in addressing these challenges. By operating directly on GLs, ngsAMOVA avoids the biases introduced by hard genotype calls and preserves information that would otherwise be lost during filtering. This leads to more accurate estimates of genetic differentiation, divergence, and phylogenetic structure across a wide range of sequencing depths and demographic scenarios. The ability to generate well-calibrated confidence intervals using genomic block bootstrapping further strengthens the method’s applicability to real-world genomic datasets, where assumptions of site independence often do not hold. Given its compatibility with standard VCF inputs and its flexibility in handling different types of hierarchical population structures, ngsAMOVA represents a practical and scalable tool for evolutionary and ecological genomics studies, particularly in cases where sequencing depth is limited or sample quality is variable.

Beyond improved accuracy, our integration of genomic block bootstrapping enables the estimation of confidence intervals for AMOVA Φ-statistics and enhances its practical utility. This approach provides a more reliable alternative to permutation-based hypothesis testing in genomic settings, where linkage disequilibrium and data dependence violate the assumptions of classical resampling schemes.

We presented ngsAMOVA, a novel GL-based framework for constructing pairwise distance matrices, performing AMOVA, *d*_*XY*_, and constructing NJ trees with low- and medium-depth NGS data. By leveraging GLs, the method circumvents limitations associated with genotype calling and enables reliable inference across a range of evolutionary scenarios. We also introduced a block bootstrapping approach for obtaining the confidence limits on the estimator based on a distribution of estimator values, and used it to quantify the significance of the results. We demonstrated the performance of ngsAMOVA through simulations. Comparing the results of this method with those obtained using genotype calls, we showed that the GL method provides more accurate results for low- and medium-depth NGS data compared to genotype calling for AMOVA, *d*_*XY*_ and Neighbor-Joining analyses, and can be used to obtain reliable estimations down to a read depth of 0.1*×*. These strengths, combined with its computational efficiency and compatibility with standard file formats, make ngsAMOVA a valuable tool for researchers analyzing population structure and genetic divergence in challenging sequencing contexts.

Nonetheless, the method assumes diploid genomes and biallelic variants and does not currently support polyploid data or multiallelic sites. Future extensions could include compatibility with polyploid data and multi-allelic variants to broaden its applicability to more complex organisms and variant types. Integrating joint inference of population structure and hierarchical grouping could further enhance the accuracy in scenarios where the hierarchical grouping labels are ambiguous or incomplete.

## Funding

This work was supported by Lundbeck Foundation Centre for Disease Evolution: [R302-2018-2155 to IA, LZ]; and; Carlsberg Foundation Young Researcher Fellowship awarded by the Carlsberg Foundation in 2019 [CF19-0712 TSK,LZ].

## Conflict of interest

none declared

## Notes

### Competing Interest Statement

The authors have declared no competing interest.

### Summary of Updates

Author affiliations updated to fix the missing address (address 1).

