## Supplementary Material for "ngsAMOVA: A Probabilistic Framework for Analysis of Molecular Variance, *d_XY_* and Neighbor-Joining Trees with Low Depth Sequencing Data"

### Supplemental content

**1 Section S1: The pairwise distance matrix (D) 4**

**4 Reproducibility and availability 39**

### Supplemental tables

1 Hierarchical analysis of molecular variance (AMOVA) with $L$ levels 23

### Supplemental figures

1 Graphical representation of the EM algorithm. . . . . . . . . . . 13
2 Percentile linkage disequilibrium decay using the model fit, with a percentile decay threshold of 95%. . . . . . . . . . . . . . . . . 26
3 Percentile linkage disequilibrium decay using the model fit, with a percentile decay threshold of 99%. . . . . . . . . . . . . . . . . 27
4 Coalescence simulation model 1 illustrating isolation after separation into regions. . . . . . . . . . . . . . . . . . . . . . . . . . . 34
5 Coalescence simulation model 2 illustrating gene flow between populations from different regions after separation. . . . . . . . . 36
6 Coalescence simulation model 3 illustrating complete isolation after separation into regions and populations. . . . . . . . . . . . 38
7 Absolute error of pairwise distances for all replicates and individual pairs. For each individual pair and replicate, absolute error was calculated within each model, method, depth, and contig size combination, based on the difference from the corresponding distance in the ground-truth data derived from contig size $10 \times 10^7$ using the ground-truth genomes. . . . . . . . . . . . . . . . . . . 40

8 RMSE of $d_{xy}$ for different $d_{xy}$ comparisons, methods, models across different depths. Colors represent different methods, and line types represent different contig sizes. Columns are different coalescence simulation models, and rows are different $d_{xy}$ comparisons. . . . . . . . . . . . . . . . . . . . . . . . . . . . . . . . 41
9 Average ratio of non-monophyletic populations vs. depth for different methods, models, and contig sizes. For each method, model, and contig size combination, the ratio of non-monophyletic populations was averaged across replicates. Colors indicate the method used to construct the trees, and lines connect the average ratio of non-monophyletic populations for different depths. Columns indicate the contig size, and rows indicate the coalescence model. . . . . . . . . . . . . . . . . . . . . . . . . . . . . . . . 42

### 1 Section S1: The pairwise distance matrix (D)

#### 1.1 Distance metric based on Identity-by-State (IBS)

**Definition 1.1** (Genotypic States and Genotype-to-Allele Mapping)**.** Consider a diploid individual $i$ with unphased genotypes at a biallelic locus. Define the allele set as $\{0, 1\}$, where 0 represents the major (ancestral) allele and 1 represents the minor (derived) allele.

Let $\mathcal{G}$ be the set of genotypic states, defined as

$$\mathcal{G} = \{0, 1, 2\}, \tag{1}$$

where each genotypic state $g_i \in \mathcal{G}$ denotes the number of minor alleles carried by individual $i$:

- $g_i = 0$: Homozygous major genotype (alleles $\{0, 0\}$),
- $g_i = 1$: Heterozygous genotype (alleles $\{0, 1\}$),
- $g_i = 2$: Homozygous minor genotype (alleles $\{1, 1\}$).

Each genotype $g_i \in \mathcal{G}$ is explicitly associated with a multiset of alleles where the genotype-to-allele mapping $A$ is defined by

$$A(0) = \{0, 0\}, \quad A(1) = \{0, 1\}, \quad A(2) = \{1, 1\}. \tag{2}$$

**Definition 1.2** (Joint Genotype Categories Matrix)**.** The joint genotype categories matrix $\mathbf{M}$ for individuals $i$ and $j$ is a $3 \times 3$ matrix defined as:

$$\mathbf{M} = \begin{array}{c} \\ g_i = 0 \\ g_i = 1 \\ g_i = 2 \end{array} \begin{array}{c} \begin{array}{ccc} g_j = 0 & g_j = 1 & g_j = 2 \end{array} \\ \begin{pmatrix} A & D & G \\ B & E & H \\ C & F & I \end{pmatrix} \end{array}, \tag{3}$$

where matrix entries are labeled from $A$ to $I$, corresponding to the combinations of $g_i$ and $g_j$, and each entry $\mathbf{M}_{g_i,g_j} := \mathbb{P}(g_i, g_j)$ represents the joint probability of observing genotypic states $g_i$ and $g_j$ in individuals $i$ and $j$, respectively. Each entry in the matrix is by definition non-negative and the matrix $\mathbf{M}$ satisfies:

$$\sum_{g_i=0}^{2} \sum_{g_j=0}^{2} \mathbf{M}_{g_i,g_j} = 1. \tag{4}$$

**Definition 1.3** (Allele Difference Function)**.** Let $\phi : \mathcal{A} \times \mathcal{A} \to \mathbb{R}_{\geq 0}$ be a function quantifying the difference between alleles. Specifically, define

$$\phi(a, b) = |a - b|, \tag{5}$$

which is a metric on the allele set $\mathcal{A}$.

**Definition 1.4** (Allele-Sampling Dissimilarity Function)**.** The allele-sampling dissimilarity function $\delta : \mathcal{G} \times \mathcal{G} \to \mathbb{R}_{\geq 0}$ quantifies the expected allelic difference when randomly sampling one allele from genotype $g_i$ and one allele from genotype $g_j$:

$$\delta(g_i, g_j) := \frac{1}{|A(g_i)||A(g_j)|} \sum_{a \in A(g_i)} \sum_{b \in A(g_j)} \phi(a, b). \tag{6}$$

In our diploid scenario, since $|A(g_i)| = |A(g_j)| = 2$, the function simplifies to

$$\delta(g_i, g_j) = \frac{1}{4} \sum_{a \in A(g_i)} \sum_{b \in A(g_j)} |a - b|. \tag{7}$$

**Lemma 1** (Explicit Form of $\delta(g_i, g_j)$)**.** *Given $\mathcal{A} = \{0, 1\}$ and the mapping $A$ defined above, we can explicitly compute:*

$$\delta(0,0) = \frac{1}{4}\Big(|0-0| + |0-0| + |0-0| + |0-0|\Big) = 0 \tag{8a}$$
$$\delta(0,1) = \frac{1}{4}\Big(|0-0| + |0-1| + |0-0| + |0-1|\Big) = \frac{1}{2} \tag{8b}$$
$$\delta(0,2) = \frac{1}{4}\Big(|0-1| + |0-1| + |0-1| + |0-1|\Big) = 1 \tag{8c}$$
$$\delta(1,0) = \frac{1}{4}\Big(|0-0| + |0-0| + |1-0| + |1-0|\Big) = \frac{1}{2} \tag{8d}$$
$$\delta(1,1) = \frac{1}{4}\Big(|0-0| + |0-1| + |1-0| + |1-1|\Big) = \frac{1}{2} \tag{8e}$$
$$\delta(1,2) = \frac{1}{4}\Big(|0-1| + |0-1| + |1-1| + |1-1|\Big) = \frac{1}{2} \tag{8f}$$
$$\delta(2,0) = \frac{1}{4}\Big(|1-0| + |1-0| + |1-0| + |1-0|\Big) = 1 \tag{8g}$$
$$\delta(2,1) = \frac{1}{4}\Big(|1-0| + |1-1| + |1-0| + |1-1|\Big) = \frac{1}{2} \tag{8h}$$
$$\delta(2,2) = \frac{1}{4}\Big(|1-1| + |1-1| + |1-1| + |1-1|\Big) = 0 \tag{8i}$$

For instance, when both individuals are heterozygous (8e), the four possible allele pairings $\{(0,0),(0,1),(1,0),(1,1)\}$ yield two unique outcomes $(0,0) = (1,1)$ and $(0,1) = (1,0)$ (a difference of 0 or 1), leading to an expected difference of $\frac{1}{2}$.

**Definition 1.5** (IBS Distance Metric)**.** The IBS distance metric $d_{ij}$ between individuals $i$ and $j$ is defined as the expected allele-sampling dissimilarity under their joint genotype probability distribution:

$$d_{ij} = \begin{cases} 0, & \text{if } i = j, \\ \displaystyle\sum_{g_i \in \mathcal{G}} \sum_{g_j \in \mathcal{G}} \delta(g_i, g_j)\, \mathbb{P}(g_i, g_j), & \text{if } i \neq j. \end{cases} \tag{9}$$

Here, $\mathbb{P}(g_i, g_j)$ denotes the joint genotype probability for genotypes $g_i$ and $g_j$ in the joint genotype categories matrix $\mathbf{M}$, as defined in 1.2. Expanding this sum and substituting the corresponding values of $\delta(g_i, g_j)$ given in equation 8 for $i \neq j$ yields

$$
\begin{aligned}
d_{ij} &= 0 \cdot \mathbb{P}(g_i = 0, g_j = 0) + \frac{1}{2} \cdot \mathbb{P}(g_i = 1, g_j = 0) + 1 \cdot \mathbb{P}(g_i = 2, g_j = 0) \\
&\quad + \frac{1}{2} \cdot \mathbb{P}(g_i = 0, g_j = 1) + \frac{1}{2} \cdot \mathbb{P}(g_i = 1, g_j = 1) + \frac{1}{2} \cdot \mathbb{P}(g_i = 2, g_j = 1) \\
&\quad + 1 \cdot \mathbb{P}(g_i = 0, g_j = 2) + \frac{1}{2} \cdot \mathbb{P}(g_i = 1, g_j = 2) + 0 \cdot \mathbb{P}(g_i = 2, g_j = 2) \\
&= \mathbb{P}(g_i = 2, g_j = 0) + \mathbb{P}(g_i = 0, g_j = 2) \\
&\quad + \frac{1}{2}\Big(\mathbb{P}(g_i = 1, g_j = 0) + \mathbb{P}(g_i = 0, g_j = 1) + \mathbb{P}(g_i = 1, g_j = 1) \\
&\qquad + \mathbb{P}(g_i = 2, g_j = 1) + \mathbb{P}(g_i = 1, g_j = 2)\Big)
\end{aligned}
\tag{10}
$$

Using the notation for the joint genotype categories matrix $\mathbf{M}$ entries $A$ through $I$, we can rewrite the distance metric as

$$
d_{ij} = C + G + \frac{B + D + E + F + H}{2}, \quad \text{for } i \neq j. \tag{11}
$$

#### 1.2 Properties of the IBS distance metric

To establish that the IBS distance metric $d_{ij}$ as defined above is a valid distance metric that is suitable for downstream analyses, we need to verify that it satisfies the standard metric axioms. Specifically, we must verify that the metric adheres to the identity of indiscernibles, non-negativity, and symmetry properties. In this subsection, we rigorously examine each of these properties in turn. Additionally, since the AMOVA framework requires the underlying distance metric to be Euclidean, we must also explicitly verify whether our IBS distance metric meets the Euclidean criterion.

##### 1.2.1 Identity of indiscernibles property

**Definition 1.6** (Identity of Indiscernibles)**.** A distance metric $d_{ij}$ satisfies the *identity of indiscernibles property* if and only if

$$
d_{ij} = 0 \iff i = j \quad \text{for all } i, j \tag{12}
$$

where $i$ and $j$ are individuals from the dataset. In other words, the distance from an individual to itself must be zero.

**Proposition 1.** *The IBS distance metric, defined as*

$$d_{ij} = \sum_{g_i \in \mathcal{G}} \sum_{g_j \in \mathcal{G}} \delta(g_i, g_j)\, I\!P(g_i, g_j) \tag{13}$$

*where $\delta(g_i, g_j)$ is the allele-sampling dissimilarity function defined in 7, satisfies the identity of indiscernibles property.*

*Proof.* To satisfy the identity of indiscernibles, we must have:

$$d_{ii} = \sum_{g_i \in \mathcal{G}} \sum_{g_i \in \mathcal{G}} \delta(g_i, g_i)\, \mathbb{P}(g_i, g_i) = 0, \tag{14}$$

for every individual $i$.

Consider the joint genotype categories matrix $\mathcal{J}$ for an individual measured against itself:

$$\mathcal{J} = \begin{array}{c} g_i = 0 \\ g_i = 1 \\ g_i = 2 \end{array} \overset{\begin{array}{ccc} g_i = 0 & g_i = 1 & g_i = 2 \end{array}}{\begin{pmatrix} A & 0 & 0 \\ 0 & E & 0 \\ 0 & 0 & I \end{pmatrix}}. \tag{15}$$

All off-diagonal probabilities vanish by definition. However, explicitly calculating the distance, we find:

$$d_{ii} = \delta(1,1)\, \mathbb{P}(g_i = 1, g_i = 1) = \frac{1}{2}E. \tag{16}$$

Here, since the term $\mathbb{P}(g_i = 1, g_i = 1) = E$ is a probability, it is strictly positive. Thus, we have:

$$d_{ii} = \frac{E}{2} > 0, \tag{17}$$

which contradicts the requirement that $d_{ii} = 0$.

Since we have shown an explicit counterexample, our initial proposition that our IBS distance metric satisfies the identity of indiscernibles property is false.

□

This contradiction arises because we are working with unphased genotypes. Therefore, to ensure mathematical consistency by explicitly satisfying the identity of indiscernibles property, we define the IBS distance metric with the explicit cases $i = j$ and $i \neq j$ as in equation 9.

##### 1.2.2 Non-negativity property

**Definition 1.7** (Non-negativity Property)**.** A distance metric satisfies the *non-negativity property* if and only if

$$d_{ij} \geq 0 \quad \text{for all } i, j. \tag{18}$$

**Proposition 2.** *The IBS distance metric defined in equation 9 satisfies the non-negativity property.*

*Proof.* By definition, the IBS-based distance metric $d_{ij}$ is expressed as a sum over probabilities multiplied by the allele-sampling dissimilarity function $\delta(g_i, g_j)$. Specifically,

$$\delta(g_i, g_j) \geq 0 \quad \text{and} \quad \mathbb{P}(g_i, g_j) \geq 0 \quad \text{for all } g_i, g_j \in \mathcal{G}. \tag{19}$$

Since both probabilities and the dissimilarity function are inherently non-negative, each term in the sum defining $d_{ij}$ is non-negative. Thus, the entire sum must be non-negative, yielding:

$$d_{ij} \geq 0 \quad \text{for all } i, j, \tag{20}$$

confirming that the IBS distance metric satisfies the non-negativity property. □

##### 1.2.3 Symmetry property

**Definition 1.8** (Symmetry Property)**.** A distance metric satisfies the *symmetry property* if and only if

$$d_{ij} = d_{ji} \quad \text{for all } i, j. \tag{21}$$

**Proposition 3.** *The IBS distance metric defined in equation 9 satisfies the symmetry property.*

*Proof.* We examine the difference explicitly:

$$\begin{aligned} d_{ij} - d_{ji} &= \sum_{g_i \in \mathcal{G}} \sum_{g_j \in \mathcal{G}} \delta(g_i, g_j)\, \mathbb{P}(g_i, g_j) - \sum_{g_j \in \mathcal{G}} \sum_{g_i \in \mathcal{G}} \delta(g_j, g_i)\, \mathbb{P}(g_j, g_i) \\ &= \sum_{g_i \in \mathcal{G}} \sum_{g_j \in \mathcal{G}} \big(\delta(g_i, g_j)\, \mathbb{P}(g_i, g_j) - \delta(g_j, g_i)\, \mathbb{P}(g_j, g_i)\big). \end{aligned} \tag{22}$$

Since the allele-sampling dissimilarity function $\delta(g_i, g_j)$ is symmetric by definition, we have:

$$\delta(g_i, g_j) = \delta(g_j, g_i) \quad \text{for all } g_i, g_j \in \mathcal{G}. \tag{23}$$

Additionally, joint probabilities are symmetric by definition of joint probability distributions:

$$\mathbb{P}(g_i, g_j) = \mathbb{P}(g_j, g_i) \quad \text{for all } g_i, g_j \in \mathcal{G}. \tag{24}$$

Thus, each term within the summation equals zero:

$$\delta(g_i, g_j)\,\mathbb{P}(g_i, g_j) - \delta(g_j, g_i)\,\mathbb{P}(g_j, g_i) = 0. \tag{25}$$

Therefore,

$$d_{ij} - d_{ji} = 0, \tag{26}$$

confirming that the IBS distance metric satisfies the symmetry property. □

###### 1.2.4 Euclidean property

We evaluated whether $d_{ij}$ satisfies the Euclidean property through simulations using EM optimization to obtain the distance matrix with genotype likelihood data. Specifically, we inspected the logs obtained from the `run_ngsAMOVA_AMOVA` rule in the Snakemake subworkflow `step5_run_ngsAMOVA_amova-nj-dxy.smk` (for the availability of the pipeline, see Section 4). The option `-doAMOVA 1` uses the `matrix_is_euclidean` function implemented in the program to assess the Euclidean nature of the resulting distance matrix. The program logs indicate whether the distance matrix meets Euclidean criteria. Analyses have demonstrated that, in certain instances, the resulting distance matrix fails to be Euclidean. to check whether the resulting distance matrix is Euclidean, and the program prints in the logs whether the resulting distance matrix is Euclidean.

The results revealed that there exist cases where the resulting distance matrix is not Euclidean. As the AMOVA framework requires Euclidean distances, we implemented and applied *Cailliez transformation* to the resulting distance matrix to ensure that the distances in the distance matrix used in AMOVA analyses are Euclidean. The implementation of the `-doAMOVA` option automatically evaluates the distance matrix for Euclidean properties and applies the Cailliez transformation if the given matrix is not already Euclidean.

#### 1.3 Expectation-Maximization (EM) optimization of the joint genotype categories matrix (M)

If genotypes are determined with high certainty and accuracy, the joint genotype categories matrix **M** can be tabulated using the observed allele counts. However, with low-depth sequencing data, it is favorable to utilize a genotype likelihood framework instead of calling genotypes.

**Definition 1.9** (Maximum Likelihood Estimation of the Joint Genotype Categories Matrix)**.** Let $G_s = (g_i, g_j)_s$ represent the combinations of latent genotypic states of individuals $i$ and $j$ at a focal site $s$. Define the parameter vector for joint genotype categories as

$$\eta = \{\eta_{(g_i,g_j)} : g_i, g_j \in \{0,1,2\}\}, \tag{27}$$

such that

$$\eta_{(g_i,g_j)} = \mathbb{P}(G_s \mid \eta). \tag{28}$$

where each entry $\eta_{(g_i,g_j)}$ represents the probability of observing genotypic states $(g_i, g_j)$ for individuals $i$ and $j$ at a focal site $s$.

Given a dataset comprising sequencing data $D$ with sites $s \in \{1, \dots, S\}$, the likelihood function is given by

$$\begin{aligned} P(D \mid \eta) &= \prod_{s=1}^{S} \mathbb{P}(D_s \mid \eta) \\ &= \prod_{s=1}^{S} \sum_{g_i,g_j \in \{0,1,2\}} \mathbb{P}(D_s \mid \eta, G_s)\, \mathbb{P}(G_s \mid \eta) \\ &= \prod_{s=1}^{S} \sum_{g_i,g_j \in \{0,1,2\}} \mathbb{P}(D_s \mid G_s)\, \eta_{(g_i,g_j)} \end{aligned} \tag{29}$$

where the term $\mathbb{P}(D_s \mid G_s)$, known as the genotype likelihood, is the likelihood of the observed data $D_s$ given the genotypic states $G_s$ at site $s$. The maximum likelihood estimate (MLE) of the parameters $\eta$ is then obtained as

$$\hat{\eta} = \arg\max_{\eta} \mathbb{P}(D \mid \eta). \tag{30}$$

As a closed-form solution to this problem is not available, the estimator $\hat{\eta}$ is typically obtained via numerical optimization techniques, such as the Expectation–Maximization (EM) algorithm (Dempster *et al.*, 1977) or Broyden –Fletcher–Goldfarb–Shanno (BFGS) algorithm (Shanno, 1970; Goldfarb, 1970; Fletcher, 1970; Broyden, 1970). In this study, we employ an EM algorithm to estimate $\hat{\eta}$, with the following Expectation (E-step) and Maximization (M-step) procedures.

##### 1.3.1 E-Step

For each locus $s$, the vector of posterior probabilities $\gamma_s^{(t)}$ is computed given the current estimate of the $\hat{\eta}^{(t)}$ and the observed data $D_s$

$$\gamma_s^{(t)} = \mathbb{P}(G_s \mid D_s, \hat{\eta}^{(t)}) = \frac{\mathbb{P}(D_s \mid G_s)\, \hat{\eta}^{(t)}}{\sum\limits_{g_i, g_j \in 0,1,2} \mathbb{P}(D_s \mid G_s)\, \hat{\eta}^{(t)}_{(g_i, g_j)}}, \tag{31}$$

where $\gamma_{(g_i,g_j)}$ represents the posterior probability that individuals $i$ and $j$ at locus $s$ have genotypic states $(g_i, g_j)$, respectively, and the superscript $(t)$ denotes the current iteration of the EM algorithm.

##### 1.3.2 M-Step

The expected joint genotype categories matrix $\gamma_s^{(t)}$ from the E-Step are used to update the estimate of the parameter set $\hat{\eta}^{(t+1)}$ for maximizing the likelihood function. The updated joint genotype categories matrix $\hat{\eta}^{(t+1)}$ is obtained by

$$\hat{\eta}^{(t+1)} = \frac{1}{S} \sum_{s=1}^{S} \gamma_s^{(t)}. \tag{32}$$

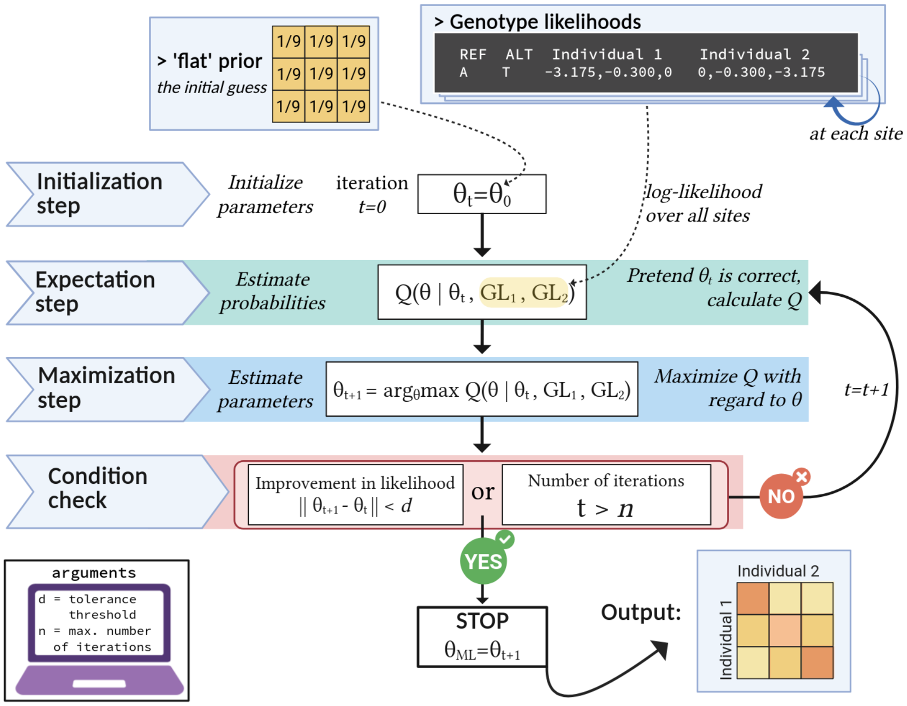

Figure S 1: Graphical representation of the EM algorithm.

### 2 Section S2: Analysis of Molecular Variance (AMOVA)

#### 2.1 Linear model of molecular variance and AMOVA

Assume a hierarchical structure with $L$ levels, where individuals are arranged into subgroups$_{(L-1)}$, the subgroups$_{(i)}$ nested within subgroups$_{(i-1)}$, and so on, until groups$_{(2)}$ nested within groups$_{(1)}$, and groups$_{(1)}$ nested within the total population. The hierarchical structure at a given level $j$ is composed of the nested arrangements of groups in all preceding levels, $G_j = G_{g_1 g_2 \ldots g_{j-1}}$, where $G_j$ is the number of groups at the $j$-th level in the hierarchical structure.

The molecular variation in this hierarchical population structure can be described using a linear model as follows:

$$\begin{aligned} X_L &= X_{g_1 g_2 \cdots g_{(L-1)} g_L} \\ &= \epsilon + x_{G_1} + x_{G_2} + \cdots + x_{G_{L-1}} + x_{G_L} \\ &= \epsilon + \sum_{j=1}^{L} x_{G_j}. \end{aligned} \tag{33}$$

where $x_j$ is the effect of the $g_j$-th subgroup within the $g_{(j-1)}$-th subgroup ($g_j = 1, 2, \ldots, G_j$), and $\epsilon$ is the unknown expectation of $X_L$. For $j = 1$, $x_{G_1}$ is the effect of the $g_1$-th group within the total population, and for $j = L$, $x_{G_L}$ is the effect of the $g_L$-th individual within the $g_{(L-1)}$-th subgroup. The effects $x_{G_1}, x_{G_2}, \ldots, x_{G_{L-1}}, x_{G_L}$ are assumed to be random and uncorrelated, and each has an associated variance component $\sigma_1^2, \sigma_2^2, \ldots, \sigma_{L-1}^2, \sigma_L^2$, respectively (Cockerham, 1969, 1973; Excoffier *et al.*, 1992; Yang, 1998).

The Analysis of Molecular Variance (AMOVA) is a statistical framework for estimating the aforementioned variance components across hierarchical subdivision levels, quantifying the differentiation within and between groups through $\Phi$-statistics, and assessing the significance of the results. The procedure of classical AMOVA analysis can be summarized in five steps:

1. Calculate the pairwise distance matrix ($\mathbf{D}$) (Section 1.1)
2. Given the hierarchical population subdivisions (Section 2.2), calculate the degrees of freedom (df; Section 2.3) and variance coefficients ($\mathbf{C}$; Section 2.4)
3. Given the distance matrix and hierarchical population subdivisions, calculate the sum of squares (SS; Section 2.5), sum of squared deviations (SSD; Section 2.6), and mean squared deviations (MSD; Section 2.7)
4. Solve for the variance components ($\sigma^2$; Section 2.8)
5. Calculate $\Phi$-statistics (Section 2.9)
6. Perform significance testing (Section 2.10)

#### 2.2 Hierarchical population subdivisions

Consider $N$ individuals arranged into a hierarchical structure with $L$ levels, as described by the command-line argument `'--formula'` with the following syntax

$$\text{Individual}_{(L)} \sim \text{Groups}_{(1)}/\text{Subgroups}_{(2)}/\ldots/\text{Subgroups}_{(L-1)} \quad (34)$$

We consider the individuals as the $L$-th level, and describe the population subdivisions at each hierarchical level $i$ by 'subgroups$_{(i)}$' for $1 < i < L-1$, 'groups$_{(1)}$' for $i = 1$ (i.e. the highest level), and 'individuals' for $i = L$ (i.e. the lowest level). The hierarchical structure can be viewed as the sum of individuals within subgroups$_{(L-1)}$, the subgroups$_{(i)}$ within subgroups$_{(i-1)}$, until the groups$_{(1)}$ within the total population.

For instance, consider a dataset of $N$ individuals, where individuals are arranged into 4 hierarchical levels, as described by the formula Individual $\sim$ Region/Population/Subpopulation. Here, region is the highest level, i.e. level 1, followed by Population (level 2), Subpopulation (level 3), and Individual (level 4) i.e. the lowest level. The total hierarchical structure in this example can be described as the sum of Individuals within Subpopulations, the Subpopulations within Populations, Populations within Regions, and Regions within the total dataset.

#### 2.3 Degrees of freedom (df)

Degrees of freedom (df) refer to the number of independent values that, given certain constraints, can vary in the calculation of a statistic. In the context of hierarchical population structures, degrees of freedom at a given level correspond to the number of independent group comparisons that can be made at a given level.

Let $k_i$ represent the number of population divisions (i.e. groups) at the $i$-th hierarchical level. The hierarchical structure at a given level $j$ is composed of the nested arrangements of groups in all preceding levels, $G_j = G_{g_1 g_2 \ldots g_{j-1}}$, where $G_j$ is the number of groups at the $j$-th level in the hierarchical structure. Then, $k_i$ can be formulated as follows:

$$
\begin{aligned}
k_0 &= 1 && \text{(35a)} \\
k_1 &= G_1 && \text{(35b)} \\
k_2 &= \sum_{g_1=1}^{G_1} G_2 && \text{(35c)} \\
k_3 &= \sum_{g_1=1}^{G_1} \sum_{g_2=1}^{G_2} G_3 && \text{(35d)} \\
&\vdots \quad \vdots\ \vdots \\
k_{L-1} &= \sum_{g_1=1}^{G_1} \sum_{g_2=1}^{G_2} \cdots \sum_{g_{L-2}=1}^{G_{L-2}} G_{L-1} && \text{(35e)} \\
k_L &= \sum_{g_1=1}^{G_1} \sum_{g_2=1}^{G_2} \cdots \sum_{g_{L-1}=1}^{G_{L-1}} G_L = N && \text{(35f)}
\end{aligned}
$$

where $k_1$ is the number of groups at the first level $G_1$ (35b); $k_2$ is the sum of the number of groups at the second level $G_2$ across all groups at the first level $G_1$ (35c); $k_3$ is the sum of the number of groups at the third level $G_3$ across all groups at the preceding levels up to the second level $G_2$ (35d); and so on, until $k_L$, sum of the number of groups (i.e. individuals, $G_L = 1$ since each individual is considered as a unique group of size 1) across all groups at all levels, equating to the total number of individuals $N$ (35f) (Yang, 1998).

The degrees of freedom for the $i$-th level within the $(i-1)$-th level, $\text{df}_i$, can be written in terms of $k$ as follows:

$$\text{df}_i = k_i - k_{i-1}. \tag{36}$$

Thus, the total degrees of freedom $\text{df}_{\text{Total}}$ is $\sum_{i=1}^{L}(k_i - k_{i-1}) = N - 1$.

#### 2.4 Variance coefficients ($c$)

Variance coefficients are used for adjusting variance component estimates to accommodate varying sample sizes within nested hierarchical structures (Weir and Cockerham, 1984; Yang, 1998).

Let $\mathbf{C} = \{c_{i,j} : 1 \leq i \leq j \leq L\}$ be an upper triangular matrix of variance coefficients, where

$$\mathbf{C} = \begin{bmatrix} c_{1,1} & c_{1,2} & c_{1,3} & \dots & c_{1,L-2} & c_{1,L-1} & c_{1,L} \\ 0 & c_{2,2} & c_{2,3} & \dots & c_{2,L-2} & c_{2,L-1} & c_{2,L} \\ 0 & 0 & c_{3,3} & \dots & c_{3,L-2} & c_{3,L-1} & c_{3,L} \\ \vdots & \vdots & \vdots & \ddots & \vdots & \vdots & \vdots \\ 0 & 0 & 0 & \dots & c_{L-2,L-2} & c_{L-2,L-1} & c_{L-2,L} \\ 0 & 0 & 0 & \dots & 0 & c_{L-1,L-1} & c_{L-1,L} \\ 0 & 0 & 0 & \dots & 0 & 0 & c_{L,L} \end{bmatrix}. \tag{37}$$

and each element $c_{i,j}$ is given by

$$c_{i,j} = \begin{Bmatrix} \frac{1}{\mathrm{df}_i}\left(v_{i,j} - \sum_{g_j=1}^{G_j} \frac{\left(N_{g_j}\right)^2}{N}\right) & \text{for } i = 1 \leq j, \\ \frac{1}{\mathrm{df}_i}\left(v_{i,j} - v_{i-1,j}\right) & \text{for } 1 < i \leq j, \\ 0, & \text{for } i > j. \end{Bmatrix}, \tag{38}$$

with $v_{i,j}$ defined as

$$v_{i,j} = \sum_{g_i=1}^{G_i} \sum_{g_j=1}^{G_j(g_i)} \frac{\left(N_{g_j}\right)^2}{N_{g_i}} \tag{39}$$

for $1 \leq i \leq j \leq L$, where $G_i$ is the number of groups at hierarchical level $i$, $G_i(g_j)$ is the number of subgroups at level $i$ contained within the $g_j$-th subgroup at level $j$, and $N_{g_i}$ is the number of individuals in the $g_i$-th subgroup at level $i$.

For $i = j$, since each $g_i$-th subgroup at level $i$ is considered as subset of itself at the same level, $G_i(g_i) = 1$, the expression simplifies to

$$\begin{aligned} v_{i,i} &= \sum_{g_i=1}^{G_i} \sum_{g_i=1}^{G_i(g_i)} \frac{\left(N_{g_i}\right)^2}{N_{g_i}} \\ &= \sum_{g_i=1}^{G_i} \frac{\left(N_{g_i}\right)^2}{N_{g_i}} \\ &= \sum_{g_i=1}^{G_i} N_{g_i} \\ &= N. \end{aligned} \tag{40}$$

For $j = L$, each $g_j$-th subgroup at level $j$ is considered as a single individual $N_{g_j} = 1$, i.e. $g_L$-th individual at level $L$, and $G_j(g_i) = N_{g_i}$, and the expression simplifies to

$$\begin{aligned} v_{i,L} &= \sum_{g_i=1}^{G_i} \sum_{g_K=1}^{G_L(g_i)} \frac{(N_{g_L})^2}{N_{g_i}} \\ &= \sum_{g_i=1}^{G_i} \sum_{g_L=1}^{N_{g_i}} \frac{(1)^2}{N_{g_i}} \\ &= \sum_{g_i=1}^{G_i} 1 \\ &= G_i. \end{aligned} \tag{41}$$

One trivial case for $c_{i,j}$ is when $i = 1$ and $j = L$, where the expression simplifies to $c_{1,L} = \frac{1}{\mathrm{df}_1}(v_{1,L} - 1)$, since $N_{g_j} = 1$ and $G_j(g_i) = N_{g_i}$ for $j = L$.

The inverse of the matrix of variance coefficients $\mathbf{C}^{-1} = \mathbf{L} = \{l_{i,j}\}$ is also an upper triangular matrix where

$$l_{i,j} = \begin{Bmatrix} \frac{1}{c_{i,i}}, & \text{if } i = j, \\ -\sum_{p=i+1}^{j} \frac{c_{i,p} l_{p,j}}{c_{i,i}}, & \text{if } i < j, \\ 0, & \text{if } i > j. \end{Bmatrix} \tag{42}$$

for $1 \leq i, j \leq L$.

#### 2.5 Sum of Squares (SS)

The conventional sum of squares is defined as the sum of squared deviations from the centroid $\bar{y}$ of a multidimensional space, where

$$\mathrm{SS} = \sum_{m=1}^{N} (y_m - \bar{y})^2. \tag{43}$$

However, calculating the centroid $\bar{y}$ may be problematic for many measures of genetic distance, as is the case with our IBS distance metric. Instead, we can express SS in terms of the normalized sum of squared pairwise Euclidean distances (Li, 1976; Anderson, 2001). This can be done due to the algebraic identity where

$$\begin{aligned}\sum_{m=1}^{N}(y_m-\bar{y})^2 &= \sum_{m=1}^{N}(y_m-\bar{y})(y_m-\bar{y}) \\ &= \sum_{m=1}^{N}(y_m-\frac{1}{N}\sum_{n=1}^{N}y_n)(y_m-\frac{1}{N}\sum_{n=1}^{N}y_n) \\ &= \sum_{m=1}^{N}\sum_{n=1}^{N}(\frac{1}{N}y_m-\frac{1}{N}y_n)(\frac{1}{N}y_m-\frac{1}{N}y_n) \\ &= \frac{1}{2N}\sum_{m=1}^{N}\sum_{n=1}^{N}(y_m-y_n)^2.\end{aligned} \tag{44}$$

Hence, the equation 43 can be rewritten as

$$\mathrm{SS} = \frac{1}{N}\sum_{1\le m<n\le N}(y_m-y_n)^2, \tag{45}$$

where the distance $y_i - y_j$ is the pairwise distance between the $i$-th and $j$-th individuals.

Let $\mathbf{D} = \{d_{m,n} : 1 \le m < n \le N\}$ be an upper triangular matrix of pairwise distances, with $d_{m,n}$ denoting the distance between the $m$-th and $n$-th individuals within the dataset. As shown in Section 1.1, our distance metric $d_{m,n}$ is symmetric, $d_{m,n} = d_{n,m}$, and the distance between an individual and itself is assumed to be zero, $d_{m,m} = 0$, for all individuals $m$ and $n$. Therefore, we can disregard the lower triangular and the diagonal of the matrix.

Let $N_{g_i}$ be the number of individuals belonging to the $g_i$-th subgroup at level $i$. We can define the submatrix $\mathbf{D}^{g_i} = \left[d_{m,n\in g_i}\right]_{N_{g_i}\times N_{g_i}}$, where $d_{m,n}$ is the distance between the $m$-th and $n$-th individuals in the $g_i$-th group at level $i$. Then, we can define the sum of squares within the $i$-th hierarchical level ($\mathrm{SS}_i^{(w)}$) as

$$\mathrm{SS}_i^{(w)} = \sum_{g_i=1}^{G_i}\frac{1}{N_{g_i}}\sum_{1\le m<n\le N_{g_i}} d_{m,n}^2 \tag{46}$$

and the total sum of squares is given by

$$\mathrm{SS}_{\mathrm{Total}} = \sum_{1\le m<n\le N}\frac{1}{N}\, d_{m,n}^2. \tag{47}$$

#### 2.6 Sum of Squared Deviations (SSD)

We can define $\mathrm{SSD}_i$, sum of squared deviations among the $i$-th level within the $(i-1)$-th level, in terms of the differences between the sum of squares as

$$\mathrm{SSD}_i = \Delta\mathrm{SS}_i = \left\{\begin{aligned} &\mathrm{SS}_{i-1}^{(w)} - \mathrm{SS}_i^{(w)}, && \text{if } 1 < i < L, \\ &\mathrm{SS}_{\mathrm{Total}} - \mathrm{SS}_i^{(w)}, && \text{if } i = 1, \\ &\mathrm{SS}_{i-1}^{(w)}, && \text{if } i = L. \end{aligned}\right\}. \tag{48}$$

Then, the total sum of squared deviations is given by $\mathrm{SSD}_{\mathrm{Total}} = \sum_{i=1}^{L} \mathrm{SSD}_i$.

#### 2.7 Mean Squared Deviations (MSD)

Let $\mathrm{MSD}_i$ be the mean squared deviations among the $i$-th level within the $(i-1)$-th level. We can define MSD as follows:

$$\mathrm{MSD}_i = \frac{\mathrm{SSD}_i}{\mathrm{df}_i} \tag{49}$$

for $1 \leq i \leq L$.

#### 2.8 Variance Components ($\sigma^2$)

Let $\mathbf{m}$ be the vector of mean squared deviations $\mathbf{m} = \{\mathrm{MSD}_i : 1 \leq i \leq L\}$. Let $\boldsymbol{\sigma^2}$ be the vector of variance components, $\boldsymbol{\sigma^2} = \{\sigma_i^2 : 1 \leq i \leq L\}$. Then, the $1 \times L$ matrix of expected mean squared deviations, $\mathbf{M}$, is given by

$$\begin{aligned} \mathbf{M} &= \mathbb{E}\left[\mathbf{m}\right] \\ &= \mathbf{C}\boldsymbol{\sigma^2}. \end{aligned} \tag{50}$$

Then, the method-of-moment estimator of the variance components $\boldsymbol{\sigma^2}$ is $\hat{\boldsymbol{\sigma^2}}$, which can be obtained from $\mathbf{M} = \mathbf{C}\hat{\boldsymbol{\sigma^2}}$ as

$$\hat{\boldsymbol{\sigma^2}} = \mathbf{C}^{-1}\mathbf{M}, \tag{51}$$

and is an unbiased estimator since

$$\begin{aligned} \mathbb{E}\left[\hat{\boldsymbol{\sigma^2}}\right] &= \mathbf{C}^{-1}\mathbf{M} \\ &= \mathbf{C}^{-1}\mathbf{C}\boldsymbol{\sigma^2} \\ &= \boldsymbol{\sigma^2}. \end{aligned} \tag{52}$$

Thus, the AMOVA estimator $\hat{\boldsymbol{\sigma}^2}$ of the variance components $\boldsymbol{\sigma}^2$ can be obtained from the expected mean squared deviations $\mathbf{M}$ as follows:

$$\hat{\sigma_i^2} = \sum_{j=i}^{L} l_{i,j}\mathbf{M}_j \tag{53}$$

for $i = 1, 2, \ldots, L$.

#### 2.9 Φ-Statistics

For an $L$-level hierarchical population structure, the Φ-statistics can be generalized in terms of the genetic correlations relative to the total variance, denoted as $\Phi_{iT}$ for $1 \leq i \leq L$, and the correlations with varying references, $\Phi_{i(i-1)}$ for $1 < i \leq L$. Disregarding the within-individual$_{(L)}$ component, we omit $\Phi_{i(i-1)}$ and $\Phi_{iT}$ for $i = L$, and formulate the Φ-statistics for any $L$-level hierarchical population structure as follows:

$$\Phi_{iT} = \sum_{j=1}^{i} \frac{\sigma^2_{j(j-1)}}{\sigma^2_{\text{Total}}}, \text{ for } 1 \leq i < L, \tag{54}$$

and

$$\Phi_{i(i-1)} = \frac{\sigma_i^2}{\sum\limits_{j=i}^{L} \sigma_j^2}, \text{ for } 1 < i < L, \tag{55}$$

where

$$\begin{aligned}(1 - \Phi_{iT}) &= \left(1 - \Phi_{i(i-1)}\right)\left(1 - \Phi_{(i-1)T}\right) \\ &= \prod_{i=1}^{L} \left(1 - \Phi_{i(i-1)}\right).\end{aligned} \tag{56}$$

To illustrate, consider a hierarchical population structure with 3 levels ($L = 3$), where individuals$_{(3)}$ are arranged into populations$_{(2)}$, and populations$_{(2)}$ are nested within regions$_{(1)}$. The Φ-statistics (including the within-individual$_{(3)}$ component) are $\Phi_{1T}$, $\Phi_{2T}$, $\Phi_{21}$, $\Phi_{3T}$, $\Phi_{32}$ (denoted as $\Phi_{CT}$, $\Phi_{ST}$, $\Phi_{SC}$, $\Phi_{IT}$, and $\Phi_{IC}$ in the classical AMOVA framework (Excoffier *et al.*, 1992), respectively), represent the genetic correlations relative to the total variance for populations$_{(2)}$, regions$_{(1)}$, and individuals$_{(3)}$, and the genetic correlations with varying references for populations$_{(2)}$ within regions$_{(1)}$, and individuals$_{(3)}$ within populations$_{(2)}$, respectively. Disregarding the within-individual$_{(3)}$ component, we omit $\Phi_{3T}$ (i.e. $\Phi_{IT}$) and $\Phi_{32}$ (i.e. $\Phi_{IC}$).

Table S 1: **Hierarchical analysis of molecular variance (AMOVA) with $L$ levels, disregarding the within-individual$_{(L)}$ component.** The sum of squares (SS) and degrees of freedom (df) for the $i$-th level within the $(i-1)$-th level, and the variance components ($\sigma^2$) for the $i$-th level, are given.

| **Source of variation** | **df** | **SSD** |
|---|---|---|
| Among groups$_{(1)}$ within total | $\mathrm{df}_1$ | $SS_{\mathrm{T}} - SS_1$ |
| Among subgroups$_{(2)}$ within groups$_{(1)}$ | $\mathrm{df}_2$ | $SS_1 - SS_2$ |
| $\vdots$ | $\vdots$ | $\vdots$ |
| Among subgroups$_{(L-1)}$ within subgroups$_{(L-2)}$ | $\mathrm{df}_{L-1}$ | $SS_{L-2} - SS_{L-1}$ |
| Among individuals$_{(L)}$ within subgroups$_{(L-1)}$ | $\mathrm{df}_L$ | $SS_L$ |
| | **Total** | $\sum_{i=1}^{L} \mathrm{df}_i = N - 1$ |

#### 2.10 Block bootstrapping

Let $m(i, k, l)$ be the metadata function that associates individuals with their respective groups, and the groups with the higher-level groups, where $i$ is the index of a group label at level $k$ (grouping$_{(k)}$), and $l$ is the query level. For example, for a hierarchical structure with individuals$_{(3)}$ nested within populations$_{(2)}$, and populations$_{(2)}$ nested within regions$_{(1)}$, the metadata function $m(i, k, l)$ for $i$, $k = 3$, and $l = 2$ gives the index of the population$_{(l=2)}$ to which the $i$-th individual$_{(k=3)}$ belongs.

The total genomic dataset can be summarized as a matrix $\mathbf{D}$ of $N$ individuals and $B$ genomic blocks,

$$\mathbf{D} = \begin{bmatrix} g_{1,1} & g_{1,2} & \cdots & g_{1,B} \\ g_{2,1} & g_{2,2} & \cdots & g_{2,B} \\ \vdots & \vdots & \ddots & \vdots \\ g_{N,1} & g_{N,2} & \cdots & g_{N,B} \end{bmatrix} \tag{57}$$

where $g_{i,j}$ represents the $j$-th genomic block of the $i$-th individual.

Let $B$ be the total number of genomic blocks, $\mathbf{G} = \{G_1, G_2, \ldots, G_B\}$. Each block, $G_i$, contains a contiguous sequence of loci identified by a start and end position as well as the chromosome number, and is defined for the species, thus remains the same for all individuals. It can be seen as an abstraction of the columns of the total genomic dataset $\mathbf{D}$.

The block configuration aims to ensure that the loci within each block are inherited together, addressing the non-independence due to linkage disequilibrium. Furthermore, to preserve the genomic correlation structure, the blocks are defined globally for all individuals and the metadata associated with the individuals is unchanged, so that $m(i, k, l)$ of $\mathbf{D}$ is the same as $m(i, k, l)$ of $\mathbf{D}_r^*$ for all $r$. Here, $\mathbf{D}_r^*$ is the $r$-th bootstrap sample consisting of $N$ individuals and $B$ genomic blocks, and the genomic block arrangements are given by $\mathbf{G}_r^* = \{G_{r,1}^*, G_{r,2}^*, \ldots, G_{r,B}^*\}$ for the $r$-th bootstrap sample.

##### 2.10.1 Identification of genomic block intervals

We identify genomic blocks assumed to be inherited independently based on linkage disequilibrium (LD) decay. For obtaining the LD estimations with simulated data, we used `ngsLD` (Fox *et al.*, 2019).

As `ngsLD` requires beagle files as input, we used `angsd` (Korneliussen *et al.*, 2014) to obtain the beagle files from the BCF files. For this, we first estimated the major and minor alleles for each site using genotype likelihoods

(`-doMajorMinor 1`) and printed the `beagle.gz` files (`-doGlf 2`) (step 1). We then obtained the positions file required by `ngsLD` using the beagle file (step 2). Lastly, we used `ngsLD` to estimate the LD between sites (step 3). For the maximum distance between SNPs to calculate LD (`--max_kb_dist`) we used the default value (100Kb). The complete pipeline is given below:

```
# step 1
angsd -doglf 2 -doMajorMinor 1 -vcf-gl FILE.bcf -out FILE

# step 2
zcat FILE.beagle.gz \
    | cut -f1 \
    | sed -e 1d -e 's/_/\t/g' \
    > ngsld_pos.tsv

# step 3
ngsLD --geno FILE.beagle.gz \
    --probs \
    --n_ind 40 \
    --n_sites $( expr $(zcat FILE.beagle.gz | wc -l) - 1) \
    --pos ngsld_pos.tsv \
    -out FILE.ld
```

We then used the script we developed to fit an LD decay model and estimate the block sizes based on percentage decay `get_block_size.R`. The script is freely available as a part of `ngsAMOVA` program.

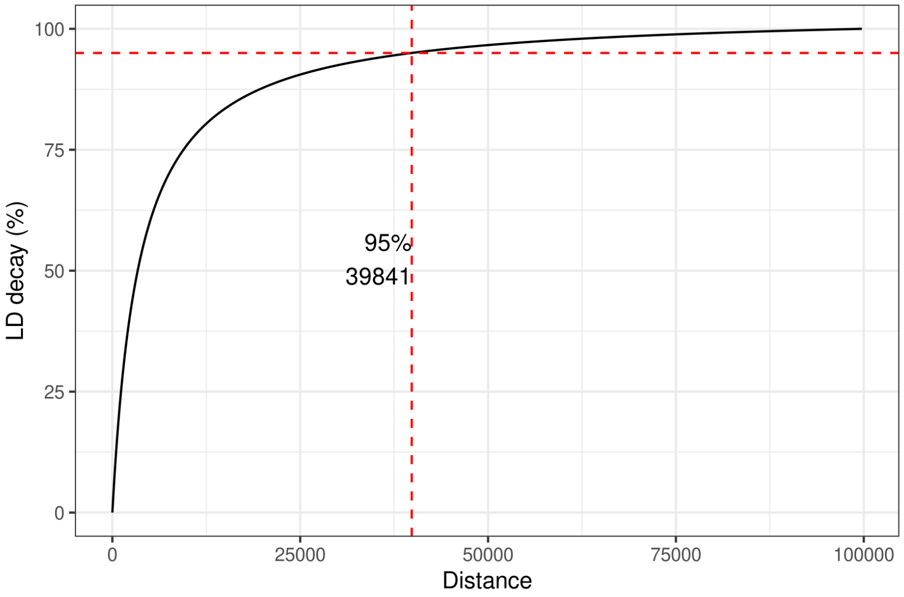

Figure S 2: Percentile linkage disequilibrium decay using the model fit, with a percentile decay threshold of 95%.

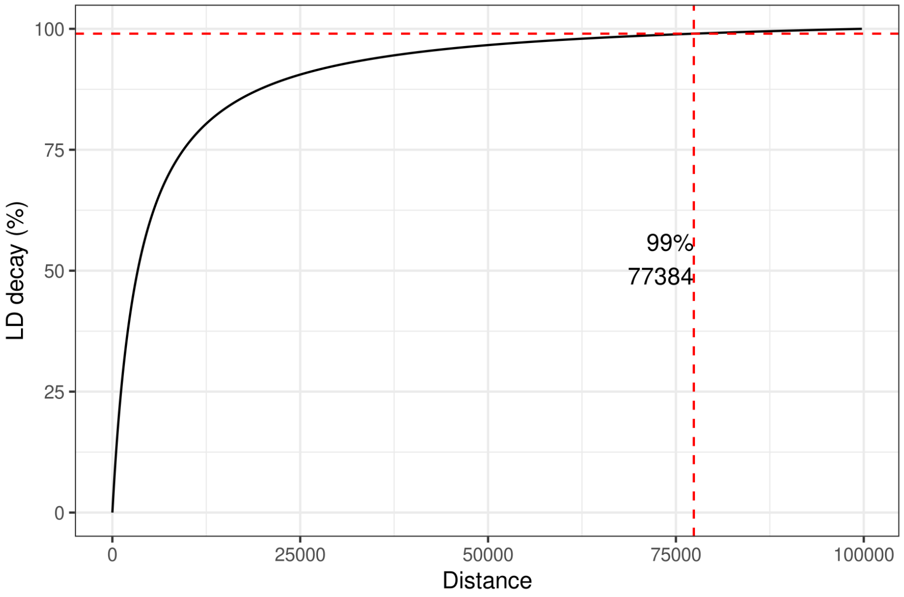

Figure S 3: Percentile linkage disequilibrium decay using the model fit, with a percentile decay threshold of 99%.

##### 2.10.2 Block bootstrapping framework

---

**Algorithm 1:** Genomic Block Bootstrapping Framework for AMOVA $\Phi$-statistic

---

**Input:** Original genomic dataset $\mathbf{D}$ consisting of $N$ individuals and $B$ blocks, the number of bootstrap replicates $R$

**Output:** Bootstrap distribution of AMOVA $\Phi$-statistic $\mathbf{\Phi}^*$

```
Φ* ← {} ;
for r = 1...R do
    G*_r ← {} ;              // initialize r-th bootstrap sample of block arrangements
    for b = 1...B do
        𝒢_b ~(i.i.d.) Unif(G) ;            // sample a block with replacement
        G*_r ← G*_r ∪ {𝒢_b} ;              // append the block to the synthetic data
    Compute the Φ-statistics Φ*_r on G*_r;
    Φ* ← Φ* ∪ φ̂*_r ;                       // append the r-th bootsrap statistic
```

---

The empirical (non-parametric) bootstrap setup for the AMOVA $\Phi$-statistic can be summarized as follows:

1. $\mathbf{D}$ is the original genomic dataset consisting of $N$ individuals and $B$ genomic blocks. $\mathbf{G} = \{G_1, G_2, \ldots, G_B\}$ genomic arrangement of blocks in $\mathbf{D}$, which is seen as a data sample drawn from a distribution $F$.

2. $\hat{\phi}$ is the $\Phi$-statistic of interest computed on the sample $\mathbf{D}$ with genomic arrangement $\mathbf{G}$ using the metadata function $m(i, k, l)$.

3. $F^*$ is the empirical distribution of the data, i.e. the resampling distribution.

4. $\mathbf{G}^* = \{G_1^*, G_2^*, \ldots, G_B^*\}$ is a bootstrap sample of the data with the same size $(B)$ as the original data

5. $\hat{\phi}^*$ is the $\Phi$-statistic of interest computed on the resample $\mathbf{D}^*$ with genomic arrangement $\mathbf{G}^*$ using the same metadata function $m(i, k, l)$ as the original computation.

After obtaining the bootstrap distribution of the $\Phi$-statistic $\mathbf{\Phi}^*$, we can estimate the confidence interval for $\hat{\phi}$. Here, we use $\hat{\phi}$ to denote the $\Phi$-statistic of interest, $\hat{\phi} \in \left(\bigcup_{i=1}^{L-1} \Phi_{iT}\right) \cup \left(\bigcup_{i=2}^{L-1} \Phi_{i(i-1)}\right)$.

###### 2.10.3 Bootstrap bias estimate

We can treat the empirical distribution, i.e. the distribution of the bootstrap estimates $\mathbf{\Phi}^* = \{\hat{\phi}_1^*, \ldots, \hat{\phi}_R^*\}$ as an approximation to the unknown distribution of $\mathbf{\Phi}$. The bias of an estimator is defined as

$$\text{Bias}(\hat{\phi}) = \mathbb{E}[\hat{\phi}] - \phi, \tag{58}$$

and the bootstrap estimate of the bias, $\widehat{\text{Bias}}_{boot}(\hat{\phi}^*, \hat{\phi})$, is given by

$$\begin{aligned} \widehat{\text{Bias}}_{boot}(\hat{\phi}, \phi) &= \bar{\phi}^* - \hat{\phi} \\ &= \frac{1}{R}\sum_{r=1}^{R} \hat{\phi}_r^* - \hat{\phi}, \end{aligned} \tag{59}$$

where $\bar{\phi}^*$ is the mean of the bootstrap estimates $\hat{\phi}_r^*$

$$\bar{\phi}^* = \frac{1}{R}\sum_{r=1}^{R} \hat{\phi}_r^*. \tag{60}$$

The bootstrap assumes the sample estimate $\hat{\phi}$ as an estimate of the population parameter $\phi$, and the bootstrap mean $\bar{\phi}^*$ as an approximation to $\mathbb{E}[\hat{\phi}]$. Here, if the center of the empirical bootstrap distribution $\bar{\phi}^*$ is at $\hat{\phi}$, then $\widehat{\text{Bias}}_{boot}(\hat{\phi}, \phi) = 0$ .

Given the bootstrap bias estimate (equation 59), we can adjust the original estimate $\hat{\phi}$ to obtain the bias-corrected estimate $\hat{\phi}_{\text{BC}}$ as

$$\begin{aligned} \hat{\phi}_{\text{BC}} &= \hat{\phi} - (\bar{\phi}^* - \hat{\phi}) \\ &= 2\hat{\phi} - \bar{\phi}^*. \end{aligned} \tag{61}$$

###### 2.10.4 Bootstrap standard error

The standard error of the bootstrap estimate $\hat{\phi}$, $\widehat{\text{SE}}_{boot}(\hat{\phi})$, is an estimate of the standard deviation of the bootstrap estimates $\hat{\phi}_r^*$. The bootstrap standard error is given by

$$\widehat{\text{SE}}_{boot}(\hat{\phi}) = \sqrt{\frac{1}{R-1}\sum_{r=1}^{R}\left(\hat{\phi}_r^* - \bar{\phi}^*\right)^2}. \tag{62}$$

##### 2.10.5 Confidence interval estimation

Confidence interval (CI) is used to estimate the likely size of a population parameter, with an associated confidence level that measures the degree of reliability of the interval. A confidence level $100(1-\alpha)\%$ implies a confidence interval that have $1-\alpha$ probability of containing the parameter. For a given population parameter $\phi$, the $100(1-\alpha)\%$ confidence interval $(\hat{\phi}_{\text{lower}}, \hat{\phi}_{\text{upper}})$ is an interval that satisfies

$$\mathbb{P}(\hat{\phi}_{\text{lower}} \leq \phi \leq \hat{\phi}_{\text{upper}}) = 1 - \alpha. \tag{63}$$

Assuming a centered interval, we have

$$P(\hat{\phi}_L \leq \phi) = P(\phi \leq \hat{\phi}_U) = \alpha/2. \tag{64}$$

We can interpret a $100(1-\alpha)\%$ confidence interval for $\phi$ as an interval that contains the true unknown parameter $\phi$ with a probability of $1-\alpha$.

**2.10.5.1 Bootstrap percentile method** The bootstrap percentile method is an empirical method that estimates the confidence interval by calculating the percentiles of the bootstrap distribution $\mathbf{\Phi}^*$. It assumes that we can use the distribution of $\mathbf{\Phi}^*$ as an approximation to the distribution of $\mathbf{\Phi}$, and that the quantiles of $\mathbf{\Phi}^*$ can be used to estimate the quantiles of $\mathbf{\Phi}$. The method uses the distribution of the bootstrap estimates $\mathbf{\Phi}^* = \{\hat{\phi}_1^*, \ldots, \hat{\phi}_R^*\}$ to form a $100(1-\alpha)\%$ confidence interval for $\hat{\phi}$:

$$\text{CI}_{(1-\alpha)} = \left(\hat{\phi}^*_{(\alpha/2)},\ \hat{\phi}^*_{(1-(\alpha/2))}\right), \tag{65}$$

where $\hat{\phi}^*_{(\alpha/2)}$ and $\hat{\phi}^*_{(1-(\alpha/2))}$ are the $100(\alpha/2)\%$ and $100(1-(\alpha/2))\%$ empirical quantiles of the bootstrap distribution $\mathbf{\Phi}^*$, respectively.

**2.10.5.2 Basic percentile method** Basic percentile (also known as reverse percentile confidence interval) differs from the percentile method in that it uses an algebraic pivot approach. It assumes that we can approximate the distributon of $\delta = \hat{\phi} - \phi$ by the distribution of $\delta^* = \hat{\phi}^* - \hat{\phi}$ where $\phi$ is the true paramater value. The calculation of the confidence intervals using the basic percentile method involves the following steps:

1. For each bootstrap sample $r$, calculate the bootstrap deviation $\delta_r^* = \hat{\phi}_r^* - \hat{\phi}$. Here, $\hat{\phi}_r^*$ is the bootstrap estimate, i.e. $\Phi$-statistic of the $r$-th bootstrap

sample, and $\hat{\phi}$ is the original estimate, i.e. the $\Phi$-statistic of the original dataset.

2. Sort the bootstrap deviations $\delta^*$ in ascending order.

3. Calculate the $100(\alpha/2)\%$ and $100(1-(\alpha/2))\%$ empirical quantiles of the bootstrap deviations $\delta^*$ to obtain $\delta^*_{(\alpha/2)}$ and $\delta^*_{(1-(\alpha/2))}$, respectively.

Here, we approximate $\delta_{(\alpha/2)}$ and $\delta_{(1-(\alpha/2))}$ by the empirical quantiles $\delta^*_{(\alpha/2)}$ and $\delta^*_{(1-(\alpha/2))}$, respectively.

**2.10.5.3 Normal approximation method** The normal approximation is a parametric method which assumes that the distribution of the $\Phi$-statistic is approximately normal, $\hat{\phi} \sim N(\phi, \sigma^2)$, and uses the normal distribution to estimate the confidence interval for $\hat{\phi}$.

If $\hat{\phi} \sim N(\phi, \sigma^2)$ with known $\sigma^2$ and unknown $\phi$, we have

$$\bar{\phi} \pm z_{\alpha/2} \frac{\sigma}{\sqrt{n}}, \tag{66}$$

and the $100(1-\alpha)\%$ CI for $\phi$ is given by

$$\mathrm{CI}_{(1-\alpha)} = \left(\bar{\phi} - z_{\alpha/2} \cdot \sigma/\sqrt{n},\ \bar{\phi} + z_{\alpha/2} \cdot \sigma/\sqrt{n}\right), \tag{67}$$

where the critical values $z$ are given by $\mathbb{P}(-z_{\alpha/2} < Z < z_{\alpha/2}) = 1-\alpha$, and $Z \sim N(0,1)$.

In the normal approximation method, we replace the unknown population standard deviation $\sigma$ with the sample standard deviation $\widehat{\mathrm{SE}}_{boot}(\hat{\phi})$ to estimate the confidence interval for $\hat{\phi}$. Then, the normal approximation $100(1-\alpha)\%$ CI for $\hat{\phi}$ can be written as

$$\mathrm{CI}_{(1-\alpha)} = \left(\hat{\phi} - z_{\alpha/2} \cdot \widehat{\mathrm{SE}}_{boot}(\hat{\phi}),\ \hat{\phi} + z_{\alpha/2} \cdot \widehat{\mathrm{SE}}_{boot}(\hat{\phi})\right). \tag{68}$$

### 3 Section S3: Simulations and Benchmarking

#### 3.1 Coalescence simulations

We used msprime (Baumdicker *et al.*, 2022) to simulate 20 replicates of three coalescence models, corresponding to 20 independent samples of individuals from the populations in the specified models . We simulated 40 diploid individuals for each model by sampling ten individuals from the four populations: *popA*, *popB*, *popC*, and *popD*. The populations in all models were distributed into two regions, namely *reg1* and *reg2*, with populations *popA* and *popB* located in region *reg1* and populations *popC* and *popD* in region *reg2*. The models were specified using demes Gower *et al.* (2022), and visualized using demesdraw Gower *et al.* (2022). Model 1 illustrates isolation after separation into regions, allowing for gene flow between populations from the same regions (Figure S4), model 2 introduces gene flow between populations from different regions (Figure S5), reducing regional isolation, and model 3 illustrates complete isolation after separation into regions and populations (Figure S6). The mutation and recombination rates were set at $1.29 \times 10^{-8}$ and $1.14856 \times 10^{-8}$, respectively. After ancestry simulations, the mutation simulations were performed using the binary mutation model to generate genetic data in VCF format. The model specifications for models 1, 2 and 3 are given in sections 3.1.1, 3.1.2 and 3.1.3, respectively.

##### 3.1.1 Model 1

```
description: model1
time_units: generations
demes:
    - name: ANC
      description: Ancestral population
      epochs:
          - {end_time: 100e3, start_size: 10000 }
    - name: reg1
      description: Region 1
      ancestors: [ANC]
      epochs:
          - {end_time: 10e3, start_size: 5000}
    - name: reg2
      description: Region 2
      ancestors: [ANC]
```

```yaml
        epochs:
            - {end_time: 10e3, start_size: 5000}
      - name: popA
        description: Population A in Region 1
        ancestors: [reg1]
        epochs:
            - {end_time: 0, start_size: 2500}
      - name: popB
        description: Population B in Region 1
        ancestors: [reg1]
        epochs:
            - {end_time: 0, start_size: 2500}
      - name: popC
        description: Population C in Region 2
        ancestors: [reg2]
        epochs:
            - {end_time: 0, start_size: 2500}
      - name: popD
        description: Population D in Region 2
        ancestors: [reg2]
        epochs:
            - {end_time: 0, start_size: 2500}
migrations:
    - {demes: [popA,popB], rate: 1e-03}
    - {demes: [popC,popD], rate: 1e-03}
    - {demes: [reg1,reg2], rate: 5e-04}
```

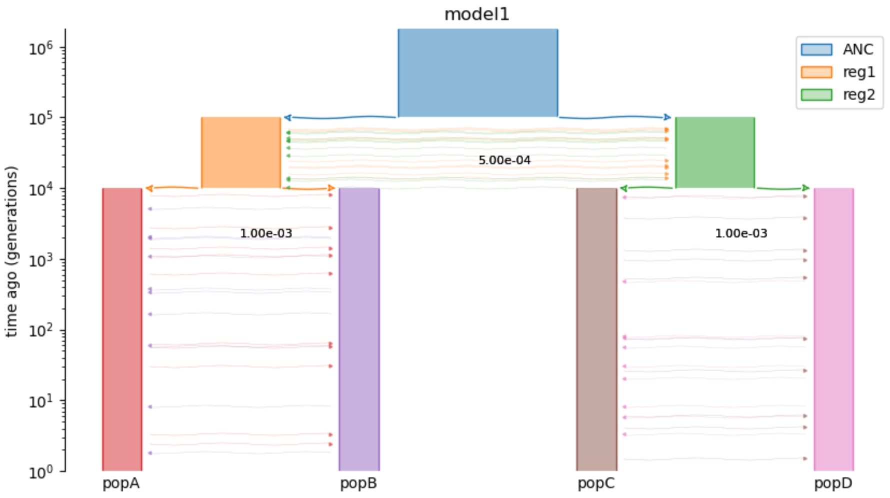

Figure S 4: Coalescence simulation model 1 illustrating isolation after separation into regions.

##### 3.1.2 Model 2

```
description: model2
time_units: generations
demes:
    - name: ANC
      description: Ancestral population
      epochs:
          - {end_time: 100e3, start_size: 10000}
    - name: reg1
      description: Region 1
      ancestors: [ANC]
      epochs:
          - {end_time: 10e3, start_size: 5000}
    - name: reg2
      description: Region 2
      ancestors: [ANC]
      epochs:
          - {end_time: 10e3, start_size: 5000}
```

```
  - name: popA
    description: Region 3
    ancestors: [reg1]
    epochs:
        - {end_time: 0, start_size: 2500}
  - name: popB
    description: Population B in Region 1
    ancestors: [reg1]
    epochs:
        - {end_time: 0, start_size: 2500}
  - name: popC
    ancestors: [reg2]
    description: Population C in Region 2
    epochs:
        - {end_time: 0, start_size: 2500}
  - name: popD
    description: Population D in Region 2
    ancestors: [reg2]
    epochs:
        - {end_time: 0, start_size: 2500}
migrations:
  - {demes: [popA,popB], rate: 1e-03}
  - {demes: [popC,popD], rate: 1e-03}
  - {demes: [popA,popC], rate: 5e-04}
  - {demes: [popA,popD], rate: 5e-04}
  - {demes: [popB,popC], rate: 5e-04}
  - {demes: [popB,popD], rate: 5e-04}
  - {demes: [reg1,reg2], rate: 5e-04}
```

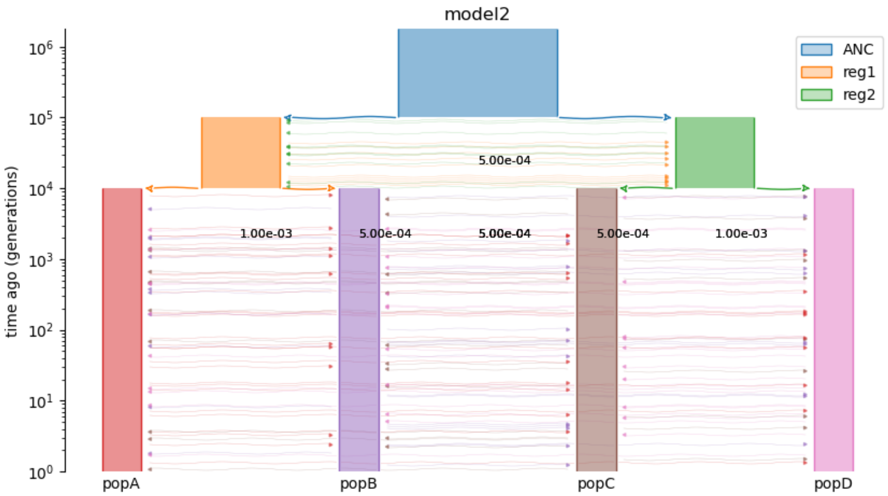

Figure S 5: Coalescence simulation model 2 illustrating gene flow between populations from different regions after separation.

##### 3.1.3 Model 3

```
description: model3
time_units: generations
demes:
    - name: ANC
      description: Ancestral population
      epochs:
          - {end_time: 100e3, start_size: 10000}
    - name: reg1
      description: Region 1
      ancestors: [ANC]
      epochs:
          - {end_time: 10e3, start_size: 5000}
    - name: reg2
      description: Region 2
      ancestors: [ANC]
      epochs:
          - {end_time: 10e3, start_size: 5000}
```

```yaml
- name: popA
  description: Population A in Region 1
  ancestors: [reg1]
  epochs:
      - {end_time: 0, start_size: 2500}
- name: popB
  description: Population B in Region 1
  ancestors: [reg1]
  epochs:
      - {end_time: 0, start_size: 2500}

- name: popC
  description: Population C in Region 2
  ancestors: [reg2]
  epochs:
      - {end_time: 0, start_size: 2500}

- name: popD
  description: Population D in Region 2
  ancestors: [reg2]
  epochs:
      - {end_time: 0, start_size: 2500}
```

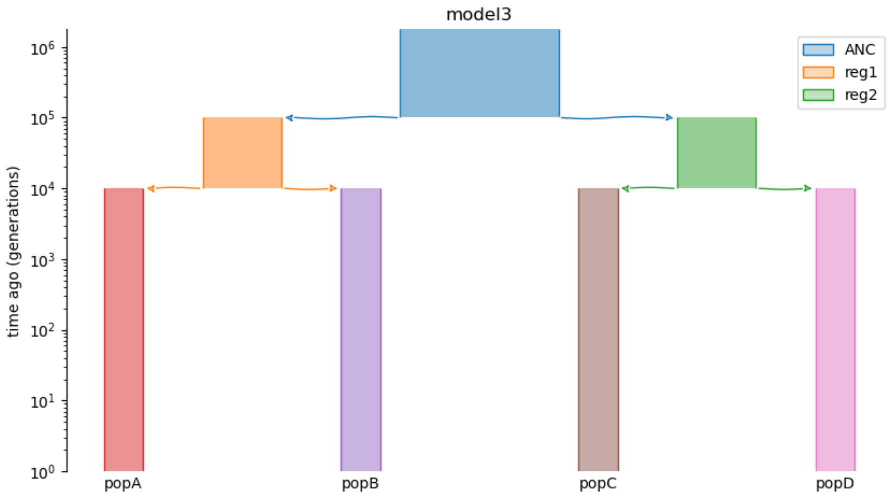

Figure S 6: Coalescence simulation model 3 illustrating complete isolation after separation into regions and populations.

#### 3.2 Sequencing data simulations

Using the data obtained from the coalescence simulations, we simulated sequencing data and genotype likelihoods with a base-calling error rate of 0.2% at varying read depths $(0.01, 0.1, 0.5, 1, 2, 5, 10, 100)$ including invariable sites (`-explode 1`) using vcfgl (Altinkaya *et al.*, 2025) with the following command line in the Snakemake pipeline:

```
({VCFGL} -i {input} \
        -o {params.prefix} \
        -O b \
        --depth {wildcards.depth} \
        --error-rate {wildcards.error_rate} \
        -addQS 1 \
        -addPL 1 \
        -explode 1 \
        -addFormatDP 1 \
        -addFormatAD 1 \
        -addInfoAD 1 \
        --rm-empty-sites 1 \
```

```
--rm-invar-sites 0 \
-doUnobserved 3 \
# GL 2: use McKenna genotype likelihood model
-GL 2 \
-printTruth 1 \
-printPileup 1 \
--seed {params.random_seed} ) 2> {log}
```

##### 3.3 Genotype calling

After sequencing data simulations, we performed genotype calling using the naive genotype calling method implemented in BCFtools (Danecek *et al.*, 2021) with the following command line:

```
bcftools +tag2tag \
        {input} -- \
        # call the GT corresponding to the most likely GL
        --GL-to-GT \
        # threshold 1: call for all non-missing GLs
        --threshold 1
```

#### 4 Reproducibility and availability

The simulation and benchmarking pipeline was developed using Snakemake (Mölder *et al.*, 2021) to ensure the reproducibility of the analyses. The pipeline is available at `https://www.github.com/isinaltinkaya/AMOVA_paper_analyses`.

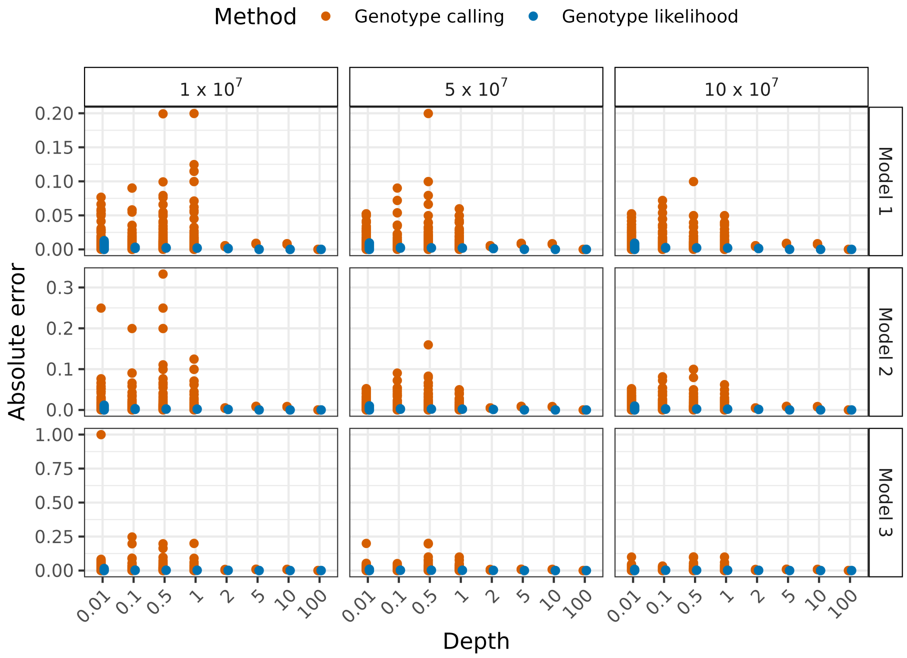

Figure S 7: Absolute error of pairwise distances for all replicates and individual pairs. For each individual pair and replicate, absolute error was calculated within each model, method, depth, and contig size combination, based on the difference from the corresponding distance in the ground-truth data derived from contig size $10 \times 10^7$ using the ground-truth genomes.

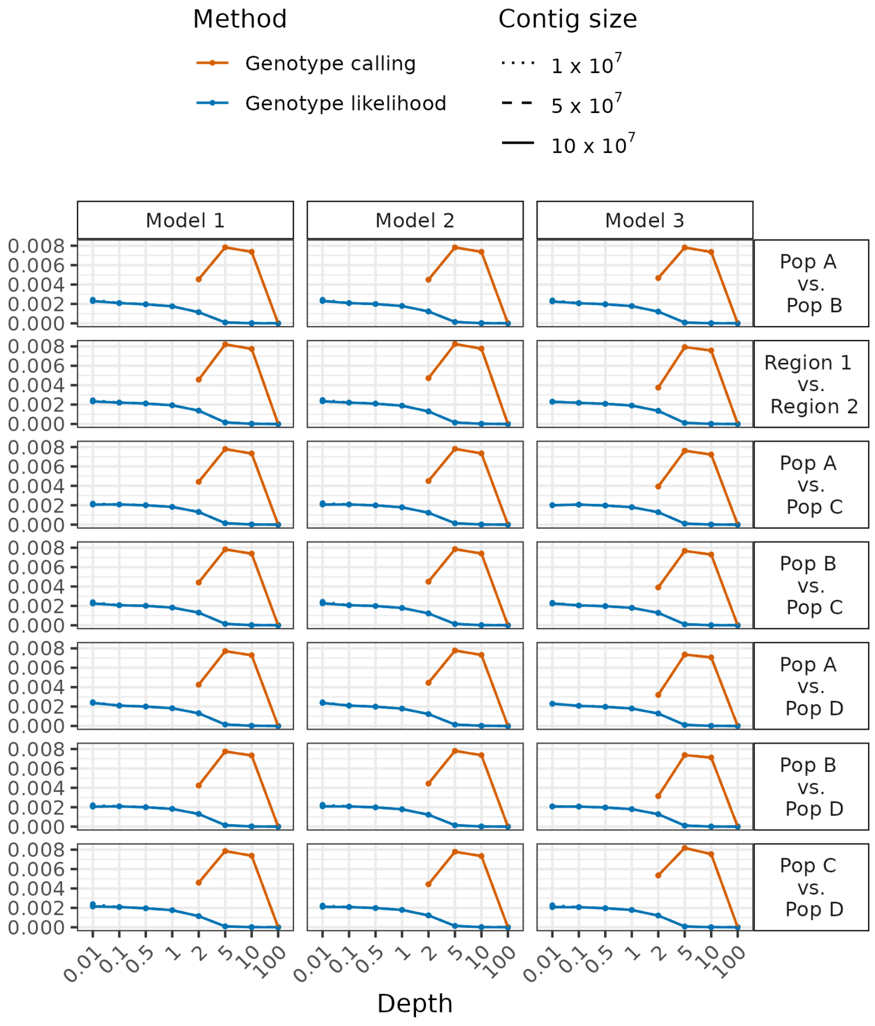

Figure S 8: RMSE of $d_{xy}$ for different $d_{xy}$ comparisons, methods, models across different depths. Colors represent different methods, and line types represent different contig sizes. Columns are different coalescence simulation models, and rows are different $d_{xy}$ comparisons.

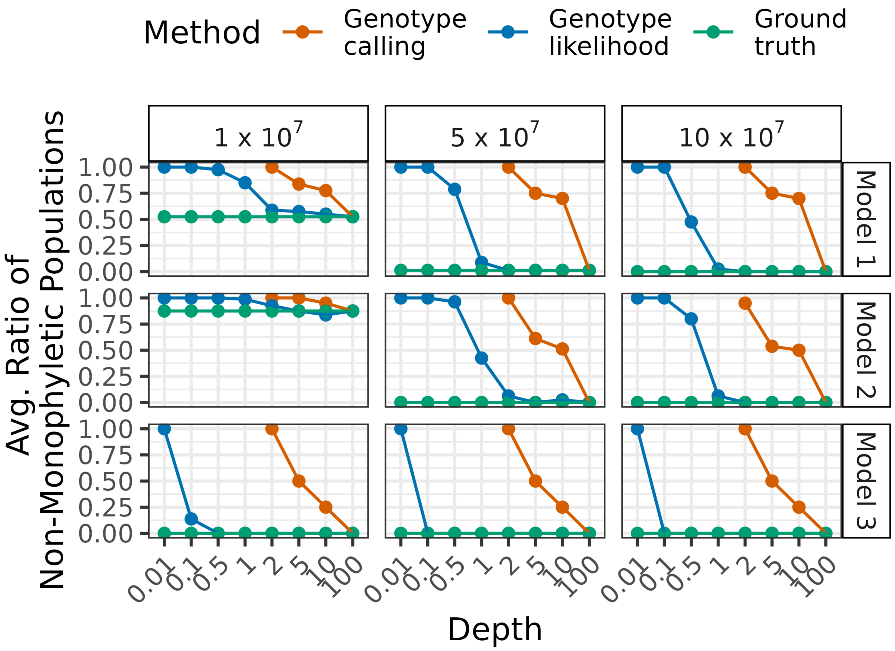

Figure S 9: Average ratio of non-monophyletic populations vs. depth for different methods, models, and contig sizes. For each method, model, and contig size combination, the ratio of non-monophyletic populations was averaged across replicates. Colors indicate the method used to construct the trees, and lines connect the average ratio of non-monophyletic populations for different depths. Columns indicate the contig size, and rows indicate the coalescence model.
